# Analytical time-dependent distributions for gene expression models with complex promoter switching mechanisms

**DOI:** 10.1101/2022.01.05.475050

**Authors:** Chen Jia, Youming Li

## Abstract

Classical gene expression models assume exponential switching time distributions between the active and inactive promoter states. However, recent experiments have shown that many genes in mammalian cells may produce non-exponential switching time distributions, implying the existence of multiple promoter states and molecular memory in the promoter switching dynamics. Here we analytically solve a gene expression model with random bursting and complex promoter switching, and derive the time-dependent distributions of the mRNA and protein copy numbers, generalizing the steady-state solution obtained in [SIAM J. Appl. Math. 72, 789-818 (2012)] and [SIAM J. Appl. Math. 79, 1007-1029 (2019)]. Using multiscale simplification techniques, we find that molecular memory has no influence on the time-dependent distribution when promoter switching is very fast or very slow, while it significantly affects the distribution when promoter switching is neither too fast nor too slow. By analyzing the dynamical phase diagram of the system, we also find that molecular memory in the inactive gene state weakens transient and stationary bimodality of the copy number distribution, while molecular memory in the active gene state enhances such bimodality.

## 1 Introduction

Gene expression in individual cells is an inherently stochastic process due to small copy numbers of biochemical molecules and probabilistic collisions between them [1]. Over the past two decades, numerous strides have been made in the models and theory of single-cell stochastic gene expression dynamics [2], which has a dual representation in terms of its probability distribution and stochastic trajectory. The former is described by a system of chemical master equations (CMEs), while the latter is described by a continuous-time Markov chain that can be simulated via Gillespie’s stochastic simulation algorithm. The simplest stochastic gene expression model is the so-called one-state model, where the promoter of the gene of interest is assumed to be always transcriptionally active [3]. This model takes into account synthesis and degradation of mRNA or protein, as well as possible transcriptional or translational bursting with the burst size having a geometric distribution, which was widely observed in experiments [4, 5]. If bursting is not considered, then the steady-state solution of the one-state model is the well-known Poisson distribution; if bursting is considered, then the steady-state solution is given by a negative binomial distribution [3], which was also extensively used in single-cell data analysis [6].

A more detailed gene expression model that is closer to the underlying molecular biology is the classical two-state model, where the promoter of the gene is assumed to switch stochastically between a transcriptionally active state and a transcriptionally inactive state [7–11]. This model has made great success in interpreting many experimental phenomena. Specifically, the two-state model can be used to reveal the biophysical origin of transcriptional bursting [12–15], which is due to a gene that is mostly inactive but transcribes a large number of mRNA molecules when it is active. Moreover, it can account for the experimentally observed bimodal distributions of mRNA and protein copy numbers that cannot be captured by the one-state model [9]. The two-state model is also analytically solvable. If bursting in each gene state is not considered, which usually occurs for the mRNA dynamics, then the steady-state distribution of the gene product number is given by a confluent hypergeometric function [7] and the time-dependent distribution can also be derived [10]. If bursting in each gene state is considered, which usually occurs for the protein dynamics, then the steady-state solution is given by a Gaussian hypergeometric function [9] and the time-dependent solution was derived in [16]. In addition, the steady-state and time-dependent solutions for the two-state model of an auto-regulatory gene network were also extensively studied [17–21].

In the two-state model, the time spent in the active or inactive gene state has an exponential distribution. This is generally a reasonable assumption for bacteria [22]. However, recent studies in mammalian cells have shown that the inactive periods for many genes may have a non-exponential peaked distribution [23, 24]. This indicates that the gene dynamics in the inactive period may contain multiple exponential steps and exhibit a “refractory” behavior: after leaving the active state, the promoter has to progress through multiple inactive states before becoming active again. In addition, there have been some experimental studies suggesting that the promoter may also have multiple active states, yielding a non-exponential active period [25–29]. In such cases, one could still use a two-state model by accepting the loss of the Markov property [30, 31]. However, adding intermediate states not only provides a convenient way to keep the Markov property, but also has the opportunity to reveal the details of the underlying biological mechanisms behind promoter switching.

In fact, gene expression models with complex multi-state promoter switching mechanisms have been widely studied from the theoretical perspective [32–39]. However, most previous papers focus on the steady-state behavior of the system, while its time-dependent behavior has received comparatively little attention. If there is only one active gene state and multiple inactive gene states, it has been found that the steady-state distribution of the mRNA copy number is given by a generalized hypergeometric function [40–44]. Thus far, there is still a lack of a detailed study about the time-dependent solution of a multi-state gene expression model with transcriptional or translational bursting. In this paper, we fill in this gap by computing the analytical time-dependent distributions of mRNA and protein copy numbers and investigate the influence of molecular memory on such distributions.

The structure of the present paper is organized as follows. In Sec. 2, we describe our multi-state model in detail and provide its CMEs. Our model can simultaneously capture non-exponential active or inactive duration distribution, as well as non-bursty or bursty gene expression pattern, and thus is very general. In Sec. 3, we obtain the analytical time-dependent solution of the multi-state model, discuss how our results reduces to previous results in one-state and two-state models, and reveal three types of possible dynamics behaviors of the system. In Sec. 4, using multiscale simplification techniques, we provide simplified expressions for the time-dependent distributions when the promoter switches rapidly or slowly between the multiple states. In Sec. 5, we investigate the phase diagram characterizing the dynamics of the multi-state model and use it to reveal the influence of molecular memory on the time-dependent distributions. We conclude in Sec. 6.

## 2 Model

The classical two-state gene expression model assumes that the promoter of a gene can switch between an active and an inactive state with the active state having a higher transcription rate than the inactive one [7]. In the two-state model, the durations of the active and inactive states both have an exponential distribution. However, recent studies have shown that in mammalian cells, the inactive periods for many genes may have a non-exponential distribution with a nonzero peak [23, 24], since the activation of the promoter is a complex multi-step biochemical process due to chromatin remodeling and the binding and release of transcription factors. This indicates that the gene dynamics in the inactive period may contain multiple exponential steps and in sum, the gene would undergo a multiple-state switching process with molecular memory (recall that an exponentially distributed transition time guarantees the Markovian property which is memoryless). Similar multi-state behavior has also been found for the active period [25–29].

Here we consider a gene expression model with complex promoter switching (Fig. 1(a)). Specifically, we assume that the promoter of the gene can exist in *L* + 1 states, denoted by *G*_0_, *G*_1_, …, *G_L_*. Biophysically, these states correspond to different conformational states during chromatin remodelling or different binding states with transcription factors [41, 42]. Moreover, we assume that the gene product of interest, i.e. mRNA or protein, is produced in a non-bursty (constitutive) or bursty manner. In each gene state *G_i_*, the synthesis of the mRNA is usually non-bursty, while the synthesis of the protein is often bursty due to rapid translation of protein from a single, short-lived mRNA molecule [13, 14]. Both non-bursty and bursty gene expression are commonly observed in naturally occurring systems [45].

In the bursty case, the synthesis of the gene product is assumed to occur in bursts of random size sampled from a geometric distribution with parameter *p*, in agreement with experiments [5]. Then the effective reactions describing the gene product dynamics are given by

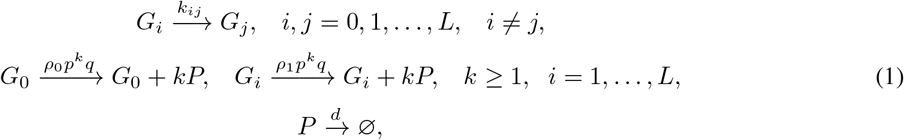

where *q* = 1 – *p*. Here the first row describes switching of the promoter between all gene states with rates *k_ij_*, the second row describes bursty production of the gene product *P* in all gene states, and the third row describes the decay of the gene product with rate *d* due to active degradation and dilution during cell division [46, 47]. Following [42], we assume that bursts occur at a rate *ρ*_0_ when the promoter is in state *G*_0_ and occur at a different rate *ρ*_1_ when the promoter is in other states *G*_1_, …, *G_L_*. If *ρ*_0_ > *ρ*_1_, then *G*_0_ is the active state and *G*_1_, …, *G_L_* are the inactive states. In this case, the active period is exponentially distributed, while the inactive period may have a non-exponential distribution. If *ρ*_0_ < *ρ*_1_, then *G*_0_ is the inactive state and *G*_1_, …, *G_L_* are the active states. In this case, the inactive period is exponentially distributed, while the active period may be non-exponentially distributed.

**Figure 1:**
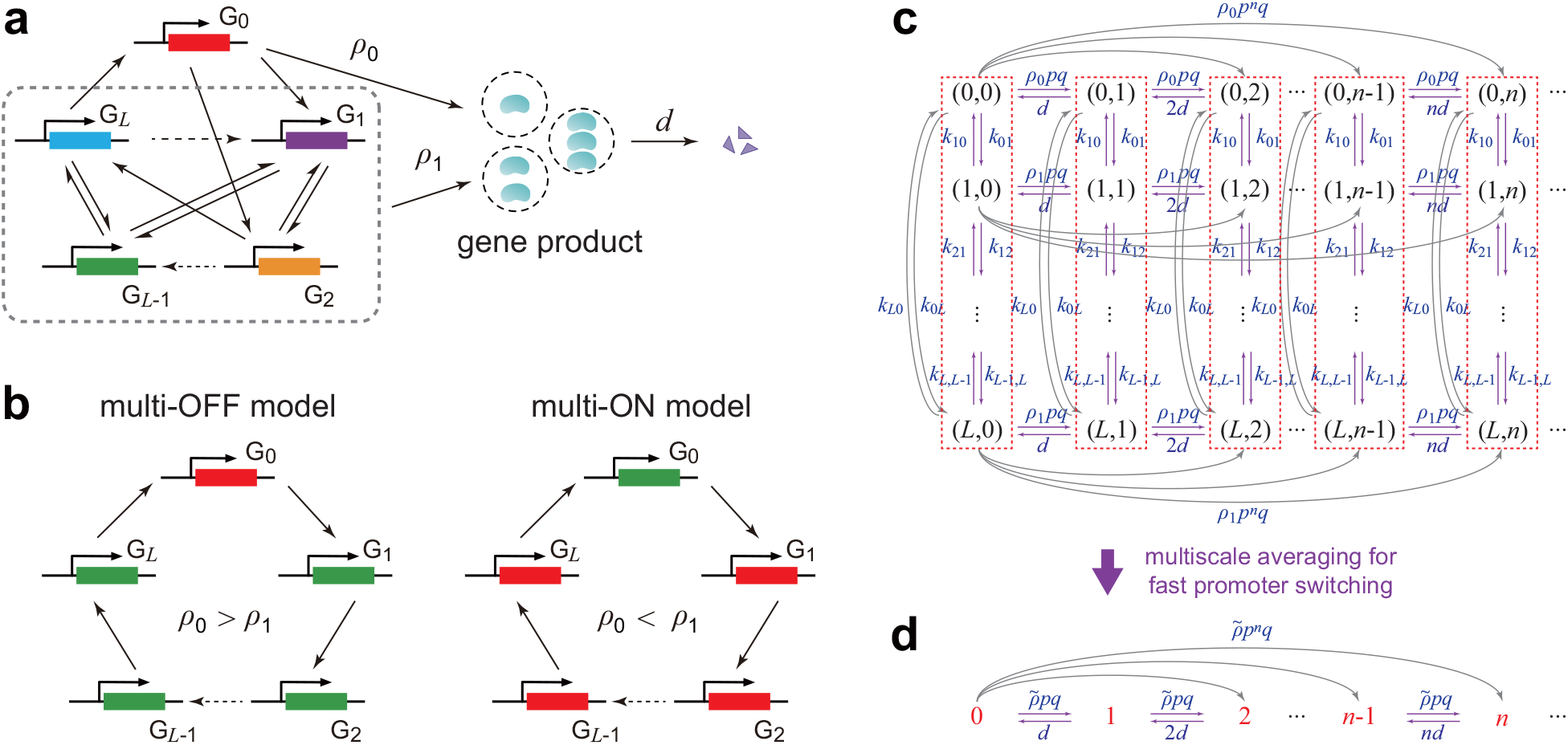
Multi-state gene expression models. (a) Schematic diagram of a multi-state model with complex promoter switching and bursting dynamics. The promoter can exist in *L* + 1 states *G*_0_, *G*_1_, …, *G_L_* and the connections between them can be arbitrary. The transcription rate is *ρ*_0_ in gene state *G*_0_ and is *ρ*_1_ in all other gene states. The gene product is produced in a bursty manner. (b) Schematic diagrams of the multi-OFF and multi-ON models, where promoter switching has an irreversible cyclic structure. For the multi-OFF model, there is only one active state and multiple inactive states. For the multi-ON model, there is only one inactive state and multiple active states. (c) Transition diagram for the Markovian dynamics associated with the multi-state model. Note that bursting can cause jumps from microstate (*i, n*) to microstate (*i, n* + *k*) for any *k* ≥ 1. This is shown for microstates (*i*, 0) in the figure but is also true for other microstates. (d) The reduced Markovian model in fast promoter conditions. When promoter switching is fast, the microstates (0, *n*), (1, *n*), …, (*L*, *n*) in (c), as enclosed by red dashed squares, are in rapid equilibrium and thus can be aggregated into a group that is labeled by group *n*.

In our model, the switching dynamics between the *L* + 1 gene states and the switching rates between them can be chosen arbitrarily, and thus the holding time in the states *G*_1_, …, *G_L_* can have a very general probability distribution. A special case occurs when promoter progression has an irreversible cyclic structure, as illustrated in Fig. 1(b) [40, 41]. If *ρ*_0_ > *ρ*_1_, then the corresponding model is referred to as the multi-OFF model since the promoter must undergo *L* inactive states one by one before entering the active state. If *ρ*_0_ < *ρ*_1_, then the corresponding model is called the multi-ON model. For both the multi-OFF and multi-ON models, the holding time in the gene states *G*_1_, …, *G_L_* is the independent sum of *L* exponential random variables and thus has a hypoexponential distribution (also called a generalized Erlang distribution).

In the non-bursty case, the effective reactions describing the gene product dynamics are given by

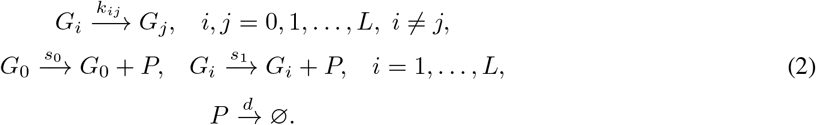

Here we assume that the gene product is produced in a constitutive manner with rate *s*_0_ when the promoter is in state *G*_0_ and is produced with a different rate *s*_1_ when the promoter is in other states *G*_1_, …, *G_L_*. Note that when the synthesis rate in the inactive gene state is zero (*s*_0_ = 0 or *s*_1_ = 0), the non-bursty model introduced above reduces to the model proposed in [42]. Interestingly, the non-bursty model described by Eq. (2) is actually a limiting case of the bursty model described by Eq. (1) [20, 48]. Since the burst size is geometrically distributed, the mean burst size is given by 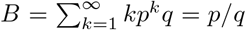. It is easy to see that when *ρ*_0_, *ρ*_1_ → ∞ and *B* → 0, while keeping *ρ*_0_*B* = *s*_0_ and *ρ*_1_*B* = *s*_1_ as constant, we have *p* → 0, *q* → 1, and

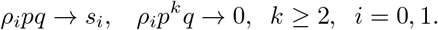

This clearly shows that the bursty model reduces to the non-bursty model in the above limit. Hence in the following, we will first derive the analytical results for the bursty model and then use them to obtain the relevant results for the non-bursty model by taking the above limit.

The microstate of the bursty model can be represented by an ordered pair (*i, n*): the state *i* of the promoter and the copy number *n* of the gene product. Let *p_i,n_*(*t*) denote the probability of having *n* copies of the gene product when the promoter is in state *i* at time *t*. Then the stochastic gene expression kinetics can be described by the continuous-time Markov chain illustrated in Fig. 1(c). The evolution of the Markovian model is governed by the following set of CMEs:

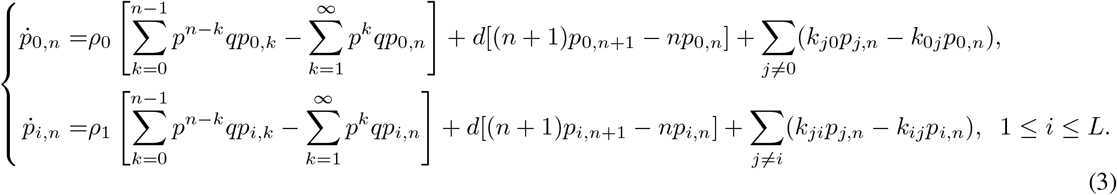

Here the first term on the right-hand side represents bursty production of the gene product, the second term represents decay of the gene product, and the last term represents promoter switching. In what follows, for convenience, we set *d* = 1 [44]. This is not an arbitrary choice but stems from the fact that the time in the CMEs can always be non-dimensionalized using the decay rate *d*. Specifically, the time given below should be understood to be non-dimensional and equal to the real time multiplied by *d*, while all other parameters *ρ_i_* and *k_ij_* given below should be understood to also be non-dimensional and equal to the real values of these parameters divided by *d*.

## 3 Analytical time-dependent solutions

### 3.1 Analytical solution for the bursty model

Before stating our results, we first introduce some notation. Let 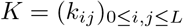 denote the generator matrix of promoter switching dynamics, where 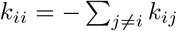 for each 0 ≤ *i* ≤ *L*. Following [44], we assume that *K* is irreducible, which means that for any pair of gene states *G_i_* and *G_j_*, there exists a transition path

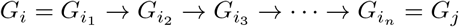

from *G_i_* to *G_j_* with positive transition rates, i.e. 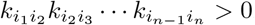. Moreover, let 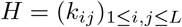 be the matrix obtained by removing the first row and the first column of the matrix *K*. Since *K* is a generator matrix, one eigenvalue of *K* must be zero. Let *μ*_1_, …, *μ_L_* denote all the nonzero eigenvalues of *−K* and let *λ*_1_, …, *λ*_*L*_ denote all the eigenvalues of *−H*. It follows from the Perron-Frobenius theorem and the irreducibility of *K* that *λ*_*i*_ and *μ_i_* all have positive real parts [44]. The physical meanings of *λ*_*i*_ and *μ_i_* are as follows. Let *p_i_*(*t*) denote the probability of being in gene state *G_i_* at time *t* and let *p*(*t*) = (*p*_0_(*t*), ⋯, *p_L_*(*t*)). Since *ṗ*(*t*) = *p*(*t*)*K*, it is clear that *μ_i_* characterize the rate for the promoter switching dynamics to relax to equilibrium. Similarly, *λ*_*i*_ characterize the rate for the gene to reach state *G*_0_ for the first time when it starts from states *G*_1_, …, *G_L_*.

To proceed, we define the following generating functions

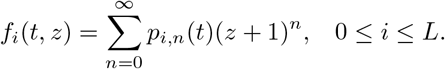

Then Eq. (3) can be converted into the following system of partial differential equations (PDEs):

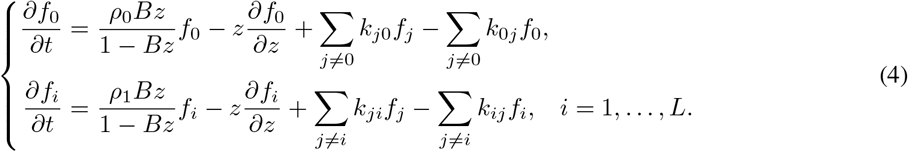

Furthermore, let 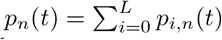 denote the probability of having *n* copies of the gene product at time *t* and let 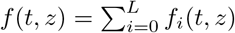 denote its generating function. For convenience, we set *ρ* = *ρ*_0_ – *ρ*_1_ and

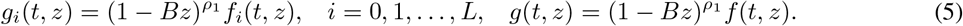

Moreover, let 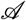 be the differential operator defined by 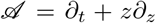. It is straightforward to check that *g_i_* satisfies the following system of PDEs:

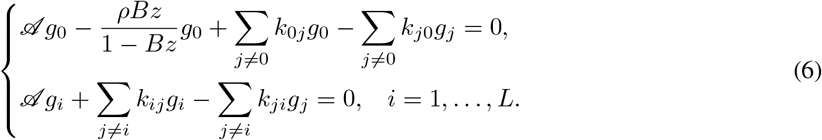

Note that Eq. (6) is much simpler than Eq. (4) since the latter contains both *ρ*_0_ and *ρ*_1_, while the former only contains their difference *ρ* = *ρ*_0_ – *ρ*_1_. Summing up all the equations in Eq. (6) yields

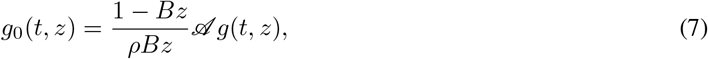

which shows that *g*_0_ can be expressed explicitly by *g*. Complex computations show that *g*_0_, *g*_1_, …, *g_L_* can all be expressed by *g* as

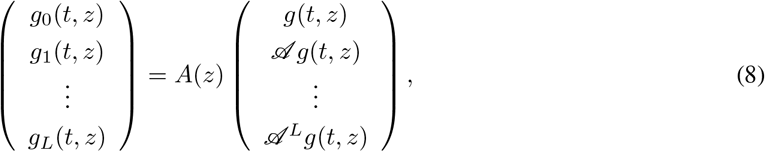

where *A*(*z*) is a square matrix of order *L* + 1 with its entries being functions of *z* (see Appendix A for the proof and the explicit expression of *A*(*z*)). Inserting Eqs. (7) and (8) into the relation 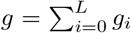, we obtain a PDE specifically for *g*, which is given by (see Appendix A for the proof)

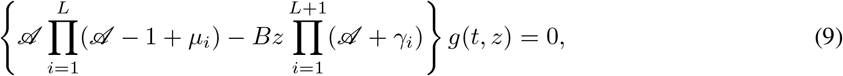

where γ_1_, …, γ_*L*+1_ are constants satisfying the equations

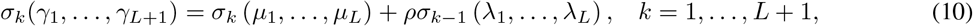

with *σ_k_*(*μ*_1_, …, *μ_L_*) being the *k*th elementary symmetric polynomials defined by

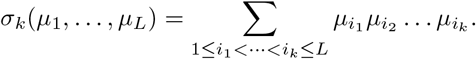

Here we assume that *σ*_0_(*μ*_1_, …, *μ_L_*) = 1 and *σ*_*L*+1_(*μ*_1_, …, *μ_L_*) = 0. Since Vieta’s formulas relate the coefficients of a polynomial to the elementary symmetric polynomials of its roots, we can see that *γ*_1_, …, *γ*_*L*+1_ are actually all the roots of a polynomial of degree *L* + 1. The fundamental theorem of algebra guarantees that *γ*_1_, …, *γ*_*L*+1_ are uniquely determined by Eq. (10).

To solve Eq. (9), we use the method proposed in [10, 16] to convert it into a solvable ordinary differential equation (ODE). To this end, we make the variable transformation *ξ* = log *z* – *t* and *η* = *z*. Let 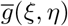 be the function with variables *ξ* and *η* that are associated with *g*(*t, z*), i.e. 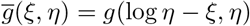. In terms of the new variables, we have

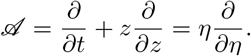

Then Eq. (9) can be converted into the ODE

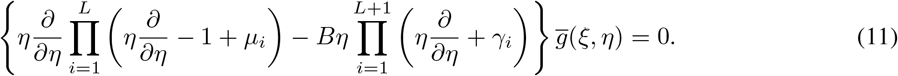

Note that for each given *ξ*, Eq. (11) is a generalized hypergeometric ODE with respect to *η* [49, Eq. 16.8.3] and its general solution is given by the linear combination of generalized hypergeometric functions [49, Eq. 16.8.6]

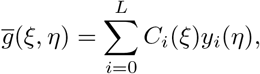

where *C_i_*(*ξ*) are constants depending on *ξ* and *y_i_*(*η*) are functions defined by

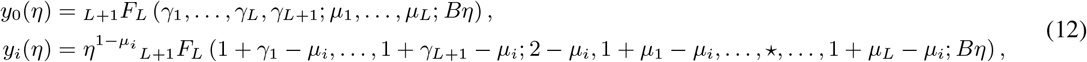

for *i* = 1, …, *L*. Here 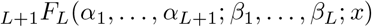 denotes the generalized hypergeometric function [49] and “★” represents that the entry 1 + *μ_i_* – *μ_i_* is removed. In terms of the original variables *t* and *z*, the function *g*(*t, z*) is given by

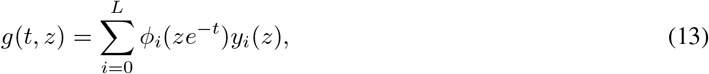

where *ϕ_i_*(*x*) = *C_i_*(log *x*).

The remaining question is to determine the functions *ϕ_i_* based on the initial conditions, or more precisely, the functions *g_i_*(0, *z*), *i* = 0, 1, …, *L* that are associated with the initial distributions. Note that for each *n* ≥ 0, it is easy to check that

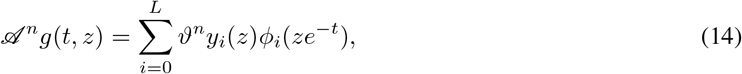

where 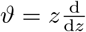. Inserting the above formula into Eq. (8) and setting *t* = 0 yield

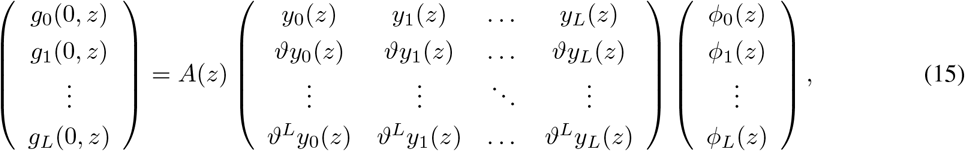

from which the functions *ϕ_i_*(*z*) can be recovered from the initial conditions *g_i_*(0, *z*). Using the transformations given in Eq. (5), in terms of the original variables *t* and *z*, the generating function *f* (*t*, *z*) is given by

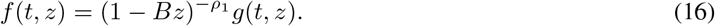

Taking partial derivatives of *f*(*t*, *z*) with respect to *z* at *z* = −1, we finally obtain the time-dependent distribution of the gene product number:

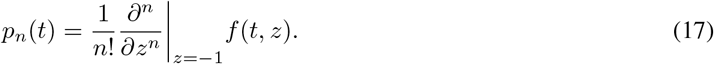

We then focus on the steady-state gene product number distribution. At the steady state, the functions *f*(*t*, *z*) and *g*(*t, z*) are independent of *t* and we denote them by *f^ss^*(*z*) and *g^ss^*(*z*), respectively. Then Eq. (9) reduces to

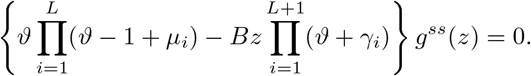

Note that this is a generalized hypergeometric ODE and its solution is given by

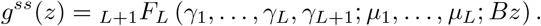

In terms of the original variable *z*, the generating function 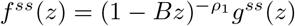, which is given by

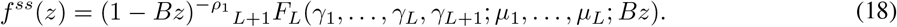

Taking derivatives on both sides of the above equation at *z* = −1 and using the differentiation formula for generalized hypergeometric functions [49, Eq. 16.3.1], we finally obtain the steady-state distribution 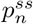 of the gene product number:

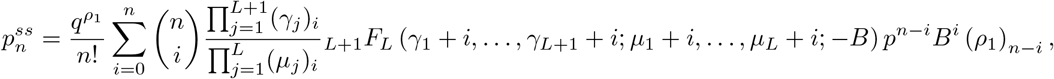

where (*x*)_*i*_ = *x*(*x* + 1) … (*x* + *i* – 1) is the Pochhammer symbol. In the special case of *ρ*_1_ = 0, i.e. no gene product molecules are produced during the inactive period, the expression can be simplified as

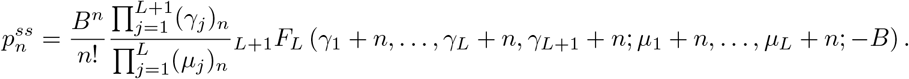

This coincides with the result obtained in [50].

### 3.2 Analytical solution for the non-bursty model

Next we focus on the non-bursty model. Recall that the non-bursty model given in Eq. (2) is a limiting case of the bursty model given in Eq. (1) when *ρ*_0_, *ρ*_1_ → ∞ and *B* → 0, while keeping *ρ*_0_*B* = *s*_0_ and *ρ*_1_*B* = *s*_1_ as constant. Under this limit, it is clear that

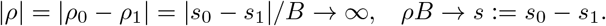

From Eq. (10), it is easy to see that *γ*_1_, …, *γ*_*L*+1_ are functions of *ρ*, *μ*_1_, …, *μ_L_*, and *λ*_1_, …, **λ*_L_*. In Supplementary Material, we have proved that

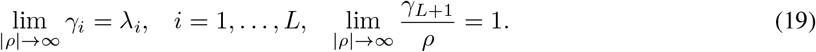

Recall that when *α_L_*_+1_ → ∞ and *z* → 0, while keeping *α*_*L*+1_*z* as constant, we have the following limit for generalized hypergeometric functions [49, Eq. 16.8.10]:

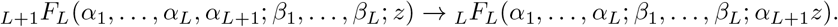

Applying this formula and taking *ρ*_0_, *ρ*_1_ → ∞ and *B* → 0, while keeping *ρ*_0_*B* = *s*_0_ and *ρ*_1_*B* = *s*_1_ as constant, we obtain the following limits:

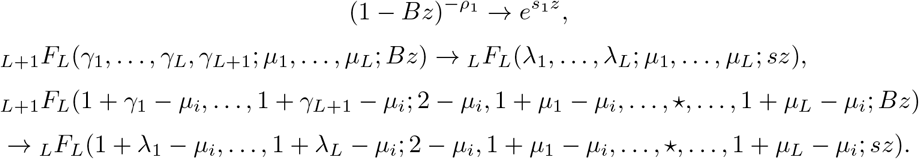

Thus from Eqs. (13) and (16), the generating function *f*(*t*, *z*) for the non-bursty model is given by

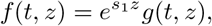

where

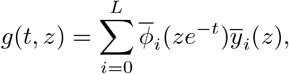

with the functions 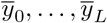 being defined by

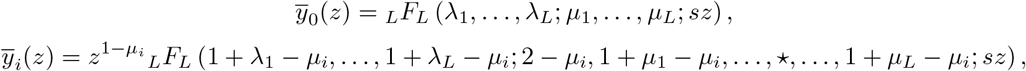

for *i* = 1, …, *L*. Here the functions 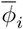 can be determined by using similar methods. Then the time-dependent copy number distribution can be recovered from *f*(*t*, *z*) based on Eq. (17).

At the steady state, it follows from Eq. (18) that the generating function reduces to

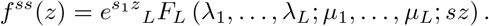

Taking derivatives on both sides of the above equation at *z* = −1 yields the steady-state copy number distribution

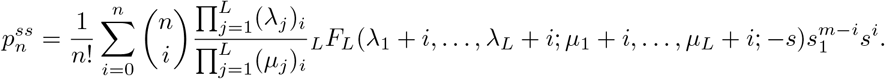

In the special case of *s*_1_ = 0, i.e. no gene product molecules are produced during the inactive period, the expression can be simplified as

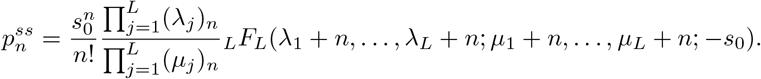

This coincides with the results obtained in [42, 44].

### 3.3 Special case of *L* = 0

We next focus on two important special cases. In the case of *L* = 0, there is only one gene state *G*_0_ and the promoter is always active. If the gene product is produced in a bursty manner, then our model reduces to the classical bursty gene expression model [3], whose effective reactions are given by

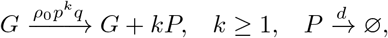

where we assume *d* = 1 for simplicity. Since *L* = 0, it follows from Eq. (13) that

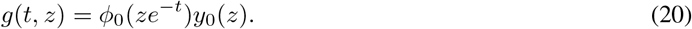

If the initial gene product number is zero, then we have *g*(0, *z*) = *f* (0, *z*) = 1. It then follows from Eq. (20) that

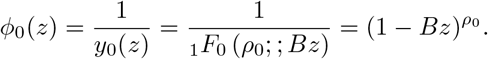

Then we obtain

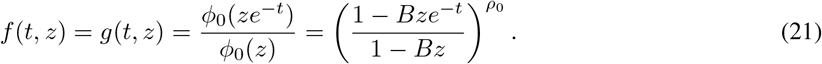

Thus when the promoter is always active, the time-dependent distribution of the gene product number can be recovered from *f*(*t*, *z*) by taking the partial derivatives with respect to *z* at *z* = −1:

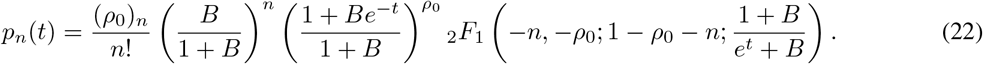

This agrees with the results obtained in [9]. Taking *t* → ∞ in the above equation yields the following steady-state distribution of the gene product number:

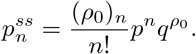

This is the well-known negative binomial distribution and agrees with the results obtained in [3].

If the gene product is produced in a non-bursty manner, then our model reduces to the following simple gene expression model:

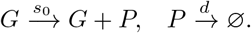

To solve this model, we only need to apply the limit *ρ*_0_ → ∞ and *B* → 0, while keeping *ρ*_0_*B* = *s*_0_ as constant, to the above results. Taking this limit in Eq. (21), we obtain

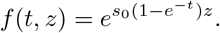

Taking the derivatives of the generating function *f*(*t*, *z*), we find that the gene product number has a Poisson time-dependent distribution that is given by

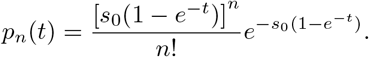

### 3.4 Special cases of *L* = 1 and *L* = 2

We next consider the special case of *L* = 1, where there are only two gene states *G*_0_ and *G*_1_. If the gene product is produced in a bursty manner, then our model reduces to the classical two-state bursty gene expression model [9, 14], whose effective reactions are given by

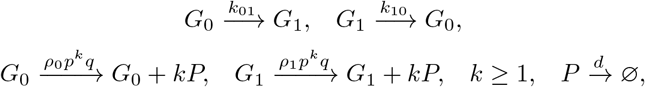

where we assume *d* = 1 for simplicity. Since *L* = 1, it is easy to check that *λ*_1_ = *k*_10_, *μ*_1_ = *k*_01_ + *k*_10_. It follows from Eq. (10) that *γ*_1_ and *γ*_2_ are determined by

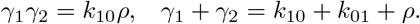

Based on Eqs. (13) and (15), we have

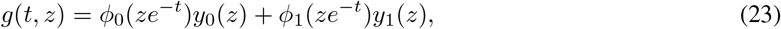

where

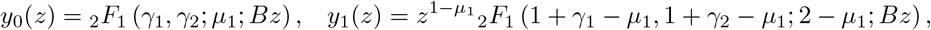

and *ϕ*_0_(*z*) and *ϕ*_1_(*z*) satisfy the following two restrictions:

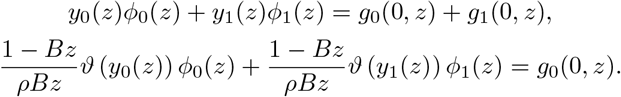

Here the first equation follows from Eq. (23) and the second one follows from Eq. (7). If the initial gene product number is zero and the promoter is in the state *G*_0_, then we have 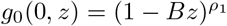 and *g*_1_(0, *z*) = 0. Solving the above two equations yield

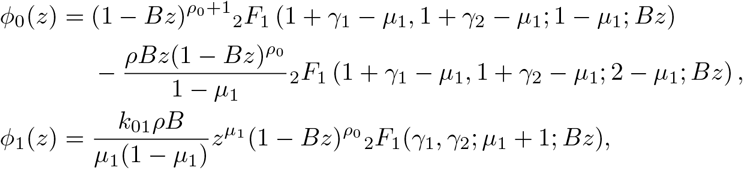

where we have used [49, Eqs. 15.10.3, 15.5.4, and 15.5.21] in the calculations. Substituting the above formulas into Eq. (23) gives the explicit expression of *g*(*t, z*). Then in terms of the original variables *t* and *z*, the generating function is given by 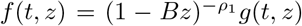. This is in agreement with the result obtained in [16]. The time-dependent copy number distribution can be recovered from *f*(*t*, *z*) by taking derivatives.

If the gene product is produced in a non-bursty manner, then our model reduces to the following classical two-state non-bursty gene expression model [7]:

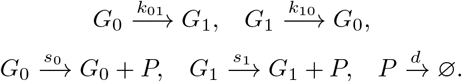

To solve this model, we consider the limit *ρ*_0_, *ρ*_1_ and *B* → 0, while keeping *ρ*_0_*B* = *s*_0_ and *ρ*_1_*B* = *s*_1_ as constant. Under this limit, it follows from Eq. (19) that *γ*_1_ → *λ*_1_ = *k*_10_ and *γ*_2_/*ρ* → 1. If the initial gene product number is zero and the promoter is in the state *G*_0_, then taking the above limit in Eq. (23) gives the generating function 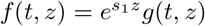, where *s* = *s*_0_ – *s*_1_ and

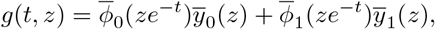

with

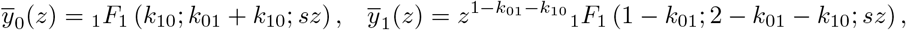

and

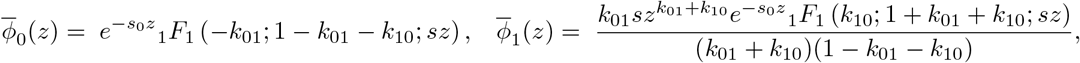

where we have used [49, Eq. 13.3.4] in the calculations. In the special case of *s*_0_ = 0, which means *G*_0_ is the inactive state and *G*_1_ is the active state, our result is consistent with the one obtained in [10].

Finally, we consider the case of *L* = 2, where the promoter can switch between three states. In this case, our model reduces to the refractory model proposed in [23]. Note that in the original refractory model, *G*_0_ is an active gene state, and the bursting in each gene state is not taken into account. The analytical time-dependent distribution for the original refractory model has been found in [51, 52]. In Appendix B, we provide the exact time-dependent solution for a three-state model with bursty production of the gene product.

### 3.5 Three types of dynamic behaviors

To validate our analytical solutions, we compare them with numerical solutions obtained using the finite-state projection algorithm (FSP) [53]. When performing FSP, we truncate the state space at a large integer *N* and solve the truncated master equation numerically using the MATLAB function ODE45. The truncation size is chosen as *N* = 5 max(*ρ*_0_*B/d, ρ*_1_*B/d*). Since *ρ*_0_*B/d* and *ρ*_1_*B/d* are the typical gene product numbers in the active and inactive gene states, respectively, the probability that the copy number is outside the truncated size is very small and practically can always be ignored. From Fig. 2, we can see that our exact solution coincides perfectly with the numerical solution obtained by the FSP at all time points.

**Figure 2:**
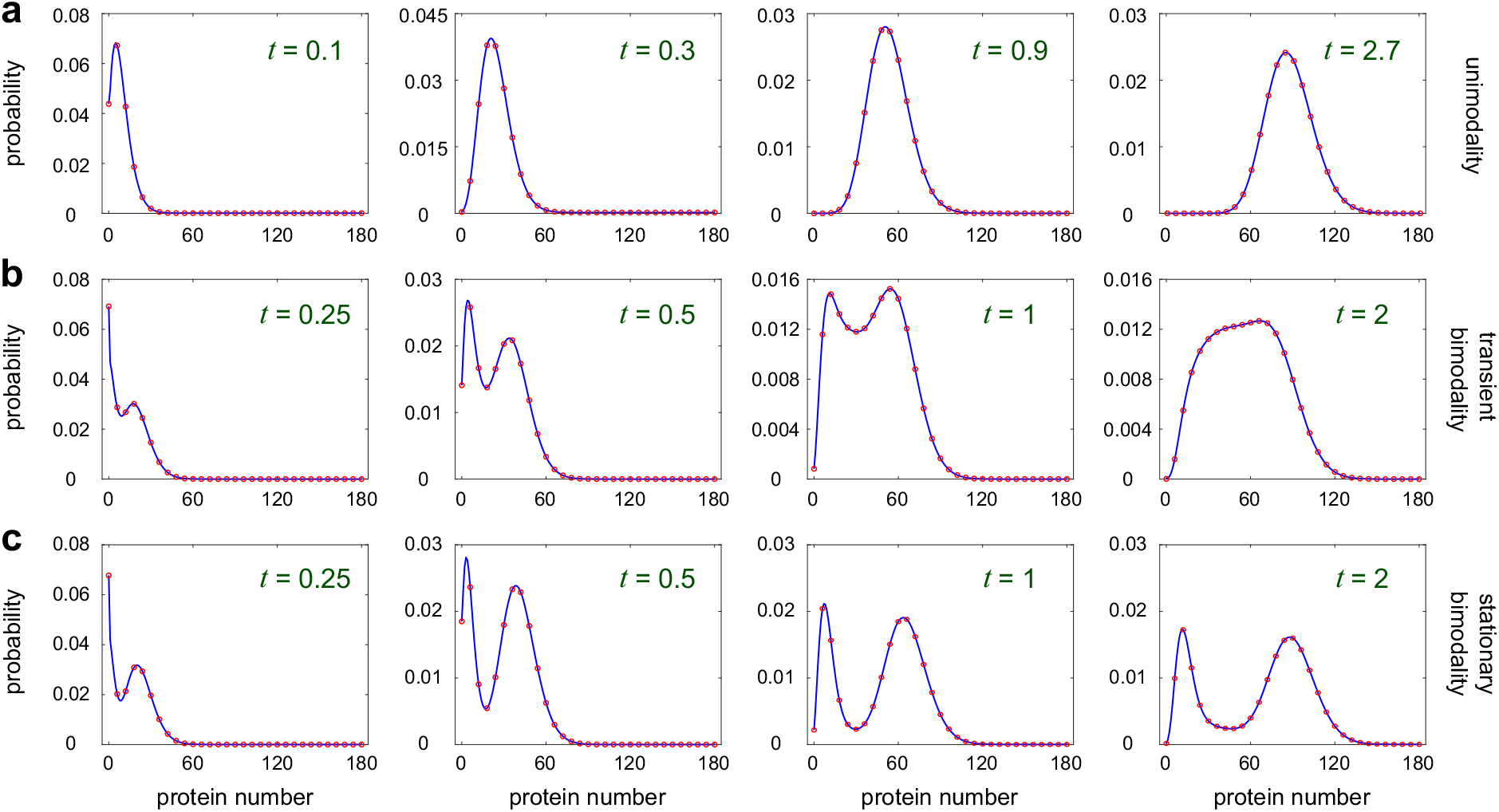
Three types of dynamic behaviors for the multi-state model. (a) Unimodality. The blue curve shows the exact solution computed via Eq. (13) and the red circles show the numerical solution computed via FSP. When computing the exact solution, we use the discrete Fourier transform algorithm proposed in [57]. This algorithm can recover the distribution directly from the generating function without calculating the higher order derivatives as in Eq. (17), which is very time-consuming and numerically unstable. Here we have assumed that *k*_01_ = *u* and *k*_12_ = *k*_23_ = … = *k*_L0_ = *Lv*. The parameters are chosen as *u* = 50, *υ* = 20, *ρ*_0_ = 240, *ρ*_1_ = 20, *B* = 1, *L* = 3. (b) Transient bimodality. The parameters are chosen as *u* = 0.65, *υ* = 1.2, *ρ*_0_ = 66, *ρ*_1_ = 10.5, *B* = 1.5, *L* = 3. (c) Stationary bimodality. The parameters are chosen as *u* = 0.1, *υ* = 0.25, *ρ*_0_ = 70, *ρ*_1_ = 10, *B* = 1.5, *L* = 3. In (a)-(c), the initial gene product number is zero and the promoter initially starts from the steady-state distribution of all gene states.

In what follows, we assume that the initial gene product number is zero and the promoter initially starts from the steady-state distribution of all gene states *G*_0_, …, *G_L_*, unless otherwise stated. The gene product number distribution can be divided into being unimodal or bimodal according to the number of modes. In general, bimodality implies multiple phenotypic states of the underlying biological system [54, 55]. To further understand the shape of the time-dependent distribution, we classify the dynamic behavior of our model into three different phases, following [21]: (i) the distribution is unimodal at all times (Fig. 2(a)); (ii) the distribution is unimodal at small and large times and becomes bimodal at intermediate times (Fig. 2(b)); and (iii) the distribution is unimodal at small times and becomes bimodal at large times (Fig. 2(c)). Multimodality with more than two modes is not detected over large swaths of the parameter space. To distinguish between them, we refer to (i) as unimodality (U), to (ii) as transient bimodality (TB), and to (iii) as stationary bimodality (SB). All the three types of dynamic behaviors can occur in our model and have been observed in naturally occurring systems (see [56, Fig. 5(c)] for U, [25, Fig. 2(c)] for TB, and [56, Fig. 2(d)] for SB).

## 4 Cases of fast and slow promoter switching

We next focus on two nontrivial special cases. The first case occurs when the promoter switches rapidly between all gene states, i.e. *k_ij_* ≫ *ρ*_0_, *ρ*_1_, *d*. In this case, the gene product number *n* is a slow variable and the promoter state *i* is a fast variable. Hence our model can be greatly simplified using the classical simplification technique of two-time-scale Markov chains called averaging [58, 59]. For simplicity, we only consider the bursty model here; the non-bursty model can be analyzed in a similar way. Since *k_ij_* are large, for each *n* ≥ 0, the microstates (0, *n*), (1, *n*), …, (*L*, *n*) are in rapid equilibrium and thus can be aggregated into a group that is labeled by group *n*, as illustrated in Fig. 1(c). In this way, our model can be simplified to the Markovian model illustrated in Fig. 1(d), whose state space is given by

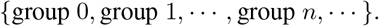

The remaining step is to compute the effective transition rates between two groups. In the fast switching regime, for each *n* ≥ 0, the microstates (*i, n*), *i* = 0, 1, …, *L* will reach a quasi-steady state. Since the transition rates between gene states are independent of *n*, the quasi-steady-state distribution is also independent of *n*. Let 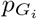 denote the quasi-steady-state probability of being in gene state *G_i_*. follows from [44, Lemma 5.3] that

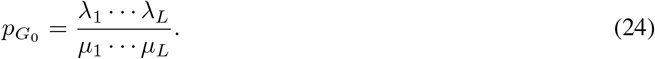

For convenience, we define the effective transcription rate as

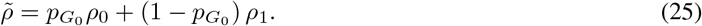

Let *q*_(*i,n*),(*i′,n′*)_ denote the transition rate from microstate (*i, n*) to microstate (*i′, n′*) for the full model. According to the averaging theory [58, 59], the effective transition rate from group *n* to group *n* + *k* is given by

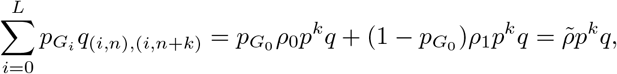

and the effective transition rate from group *n* to group *n −* 1 is given by

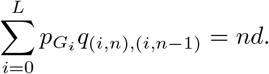

Thus far, we have obtained all effective transition rates for the group dynamics (Fig. 1(d)). Note that this is equivalent to a one-state model in the case of *L* = 0, where the transcription rate *ρ*_0_ in the gene state *G*_0_ is replaced by the effective transcription rate 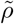. Since we have assumed that the promoter initially starts from the quasi-steady-state distribution of all gene states, the averaging theory guarantees that distributions of the full and reduced models agree with each other over the whole time axis [60]. If the initial gene product number is zero, then it follows from Eq. (22) that the approximate time-dependent distribution of the gene product number in the fast switching regime is given by

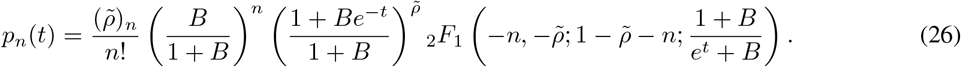

To validate this result, we compare the approximate solution with the exact solution in Fig. 3(a). It can be seen that they coincide with each other perfectly at all time points.

**Figure 3:**
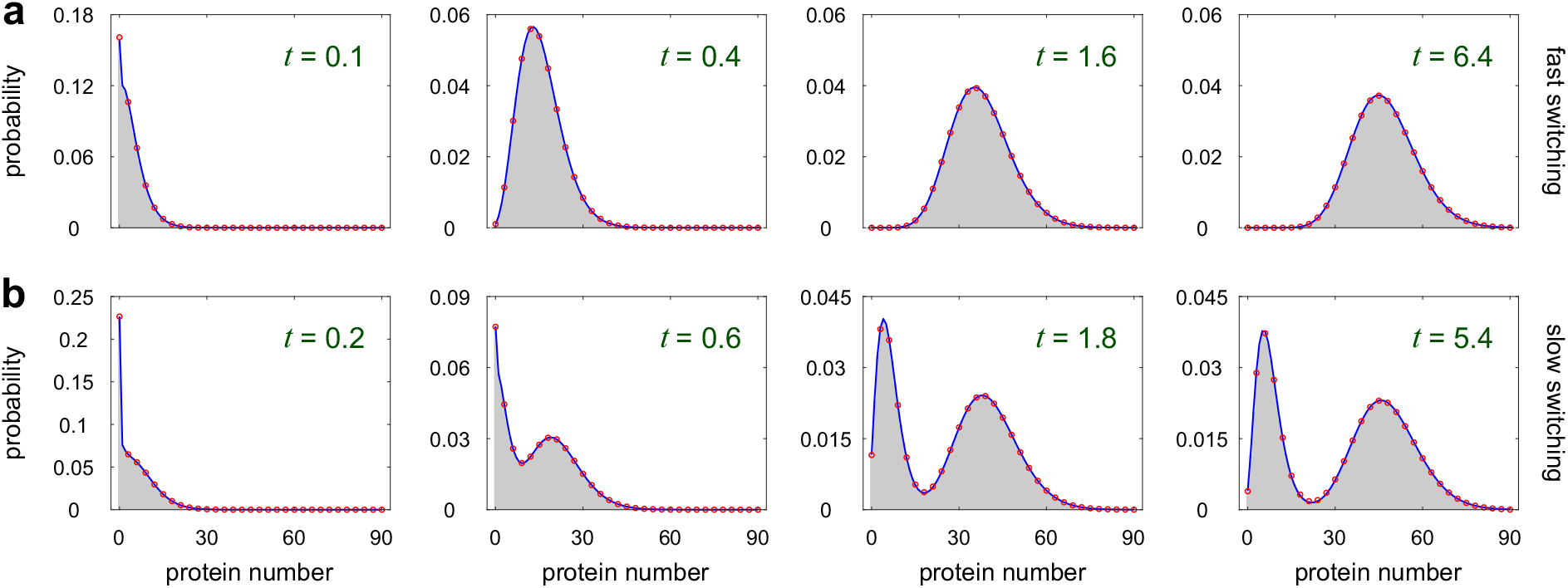
Comparison between the exact and approximate time-dependent distributions in fast and slow switching conditions for the multi-OFF model. (a) Fast promoter switching. The parameters are chosen as *u* = 100, *υ* = 200, *ρ*_0_ = 42, *ρ*_1_ = 10, *B* = 1.5. (b) Slow promoter switching. The parameters are chosen as *u* = 0.003, *υ* = 0.005, *ρ*_0_ = 32, *ρ*_1_ = 5, *B* = 1.5. In (a),(b), the blue curve shows the exact solution when *L* = 2, the red circles show the approximate solution when *L* = 2, and the grey region shows the exact solutions when *L* = 10. The exact solution is computed via Eq. (16), while the approximate solution is computed via Eq. (26) for fast switching and via Eq. (28) for slow switching. Clearly, molecular memory has no effect on the distributions in fast and slow switching conditions.

The second case occurs when the promoter switches slowly between all gene states, i.e. *k_ij_* ≪ *ρ*_0_, *ρ*_1_, *d*. In this case, the gene product number *n* is a fast variable and the promoter state *i* is a slow variable. Since *k_ij_* are small, for each 0 ≤ i ≤ *L*, the microstates (*i, n*), *n* ≥ 0 will relax to a quasi-steady state before promoter switching takes places. If the initial gene product number is zero, then it follows from Eq. (22) that the distribution *p*_*n*|*i*_(*t*) of the gene product number conditioned on gene state *G_i_* is given by

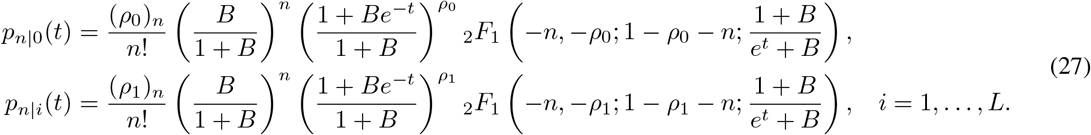

Let 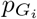 denote the steady-state probability of being in gene state *G_i_*, respectively. It follows from Eq. (24) that

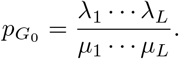

Since we have assumed that the promoter initially starts from the steady-state distribution of all gene states, the approximate time-dependent distribution of the gene product number in the slow switching regime is given by

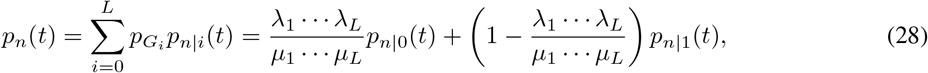

where *p*_*n*|0_(*t*) and *p*_*n*|1_(*t*) are given in Eq. (27). Taking *t* → ∞ in the above equation yields the steady-state distribution of the gene product number, which is given by

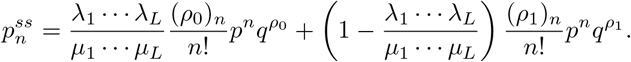

This is the mixture of two negative binomial distributions. Since a negative binomial distribution is unimodal, the mixture of two negative binomials can yield a bimodal copy number distribution. To validate this result, we compare the approximate solution with the exact solution in Fig. 3(b), from which we see that they coincide with each other perfectly at all time points.

## 5 Influence of molecular memory on the time-dependent distribution

### 5.1 Fast and slow promoter switching

The major difference between our model and the classical one-state or two-state model is the complex promoter switching mechanism, which leads to complex non-exponential distribution of the active or inactive period, i.e. molecular memory. Here we investigate the effect of molecular memory on the time-dependent distribution of the gene product number.

To do this, we focus on the multi-ON and multi-OFF models with *L* + 1 gene states (Fig. 1(b)). For the multi-ON (multi-OFF) model, the active (inactive) period has a non-exponential distribution. Here the parameter *L* can be understood as the strength of molecular memory. A larger *L* usually gives rise to a larger deviation from the exponential distribution. To make an equitable comparison between models with different values of *L*, we assume that the promoter switching rates are given by

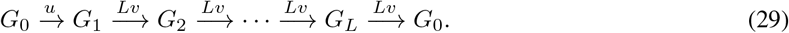

In other words, we assume *k*_01_ = *u* and *k*_12_ = *k*_23_ = … = *k*_*L*0_ = *Lv*. Under this assumption, the transition time from *G*_0_ to *G*_1_, denoted by *T*_0_, has an exponential distribution with mean 1/*u*. Moreover, the transition time from *G*_1_ to *G*_0_, denoted by *T*_1_, has an Erlang distribution with shape parameter *L* and mean 1/*v*. Therefore, the parameter *v* can be interpreted as the effective switching rate from *G*_1_ to *G*_0_. Note that the means of the two transition times are independent of *L*. The noise, characterized by the coefficient of variation squared, in *T*_1_ is given by 1/*L*. As *L* → ∞, the noise in *T*_1_ tends to zero and thus *T*_1_ converges to a point mass at 1/*v*, in which case molecular memory is the strongest.

There are two reasons for assuming Eq. (29). First, if we assume arbitrary switching rates between all gene states, then the transition time *T*_1_ may depend on many parameters and thus it is very difficult to study the influence of molecular memory on the copy number distribution. Second, in experiments, the non-exponential switching time distribution usually has a non-zero mode and is right-skewed [23]; this kind of distribution in general can be well approximated by an Erlang distribution.

To understand the influence of molecular memory on the time-dependent distribution, we first focus on the cases of fast or slow promoter switching. For both the multi-ON and multi-OFF models, the steady-state probability of being in gene state *G*_0_ is given by

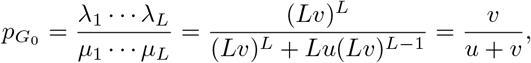

which does not depend on *L*. Since the mean holding time in gene state *G*_0_ is given by 〈*T*〉_0_ = 1/*u* and the mean holding time in gene states *G*_1_, …, *G_L_* is given by 〈*T*〉_1_ = 1/*v*, we clearly have

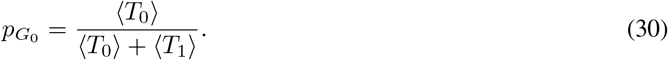

In fact, this equation holds generally for any promoter switching mechanism (Fig. 1(a)) and does not require *T*_1_ to be Erlang distributed. To see this, recall that for a Markov chain system, the mean holding time in a subset *S* of the state space is the total probability in the subset *S* divided by the total probability flux between the subset *S* and its complement *S^c^* (see [61, Theorem 1]). For any promoter switching mechanism, the total probability flux *J* between gene state *G*_0_ and other gene states *G*_1_, …, *G_L_* is given by 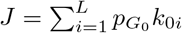. Hence the mean holding time in gene state *G*_0_ is 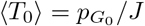 and the mean holding time in gene states *G*_1_, …, *G_L_* is 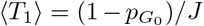. This gives the proof of Eq. (30). In the fast switching regime, the effective transcription rate given in Eq. (25) and the time-dependent distribution given in Eq. (26) are both independent of *L*. On the other hand, in the slow switching regime, the time-dependent distribution given in Eq. (28) is also independent of *L*. This shows that molecular memory has no effect on the time-dependent distribution in the limiting cases of fast and slow promoter switching, as illustrated in Fig. 3.

### 5.2 General case

We have seen that molecular memory has no influence on the time-dependent distribution when the promoter switches rapidly or slowly between all gene states. A natural question is whether molecular memory will affect the time-dependent distribution when promoter switching is neither too fast nor too slow. To see this, recall that the gene expression system can exhibit three types of dynamical behaviors: unimodality (U), transient bimodality (TB), and stationary bimodality (SB).

To understand the influence of molecular memory on the copy number distribution, we illustrate the *u* − *v* phase diagrams for the multi-ON and multi-OFF models under different choices of *L* in Fig. 4. Clearly, for both models, U occurs when promoter switching is very fast, while TB and SB can only occur when promoter switching is relatively slow. This is in sharp contrast to a positive or negative feedback genetic loop, where TB can also occur in fast switching conditions [21]. Compared with TB, the occurrence of SB requires slower promoter switching rates. Previous studies [42] have shown that for both models, the steady-state noise in the gene product number will decrease as the strength *L* of molecular memory increases. This is because the increase of *L* leads to a smaller noise in the transition time from *G*_1_ to *G*_0_ and thus will also reduce noise in the steady-state distribution. Generally, unimodal distributions have smaller noise than bimodal ones. Hence intuitively, we expect that the increase of *L* should reduce the regions of TB and SB in the phase diagram. For the multi-OFF model, we find that this is indeed the case, where the regions of TB and SB both shrink remarkably as *L* increases (Fig. 4(a)). However, for the multi-ON model, we find that the intuition is incorrect and the opposite is true — the regions of TB and SB both enlarge with the increase of *L* (Fig. 4(b)).

**Figure 4:**
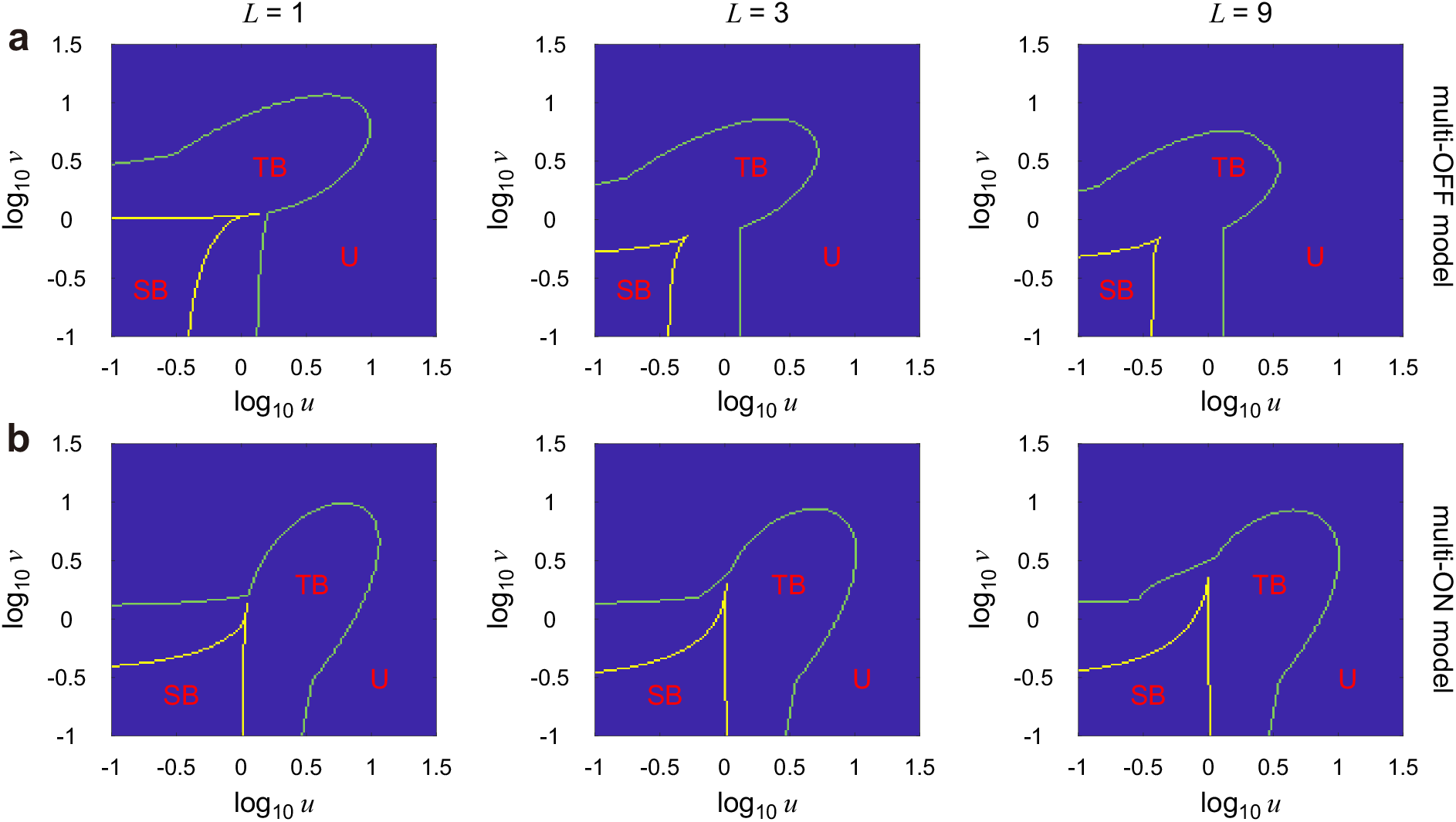
Dynamic phase diagrams for the multi-OFF and multi-ON models as *u* and *υ* vary. (a) Phase diagram for the multi-OFF model under different choices of *L*. The parameters are chosen as *ρ*_0_ = 40, *ρ*_1_ = 0, *p* = 0.3. (b) Same as (a) but for the multi-ON model. The parameters are chosen as *ρ*_0_ = 0, *ρ*_1_ = 40, *p* = 0.3. The green and yellow curves enclose the regions for TB and SB, respectively. The two curves are determined numerically as follows. First, for any bimodal distribution, we define the strength of bimodality [20] as *κ* = (*H*_low_ − *H*_valley_)/*H*_high_, where *H*_low_ is the height of the lower peak, *H*_high_ is the height of the higher peak, and *H*_valley_ is the height of the valley between them. For any unimodal distribution, *κ* is automatically set to be 0. Clearly, *κ* is a quantity between 0 and 1. The larger the value of *κ*, the stronger bimodality is. Next, we compute the value of *κ* in steady-state conditions. If *κ* ≥ 10^−6^, then the dynamical behavior is classified into SB. Otherwise, we compute the values of *κ* at a sequence of time points 0 = *t*_0_ < *t*_1_ < ⋯ < *t_N_* = *T*, where the interval between these time points is chosen to be small enough, and the final time *T* is chosen to be large enough to guarantee that the system has reached the steady state. If *κ* > 10^−6^ for some intermediate time point *t_i_*, then the dynamical behavior is classified into TB; otherwise, it is classified into U. Here the threshold 10^−6^ is chosen to be very small but non-zero in order to ensure the algorithm to be numerically stable. Both the steady-state and time-dependent distributions are computed using FSP.

To explain this counteractive phenomenon, we illustrate the steady-state copy number distribution under different choices of *L* in Fig. 5 for both the multi-OFF and multi-ON models. In Fig. 5(a), the steady-state distribution is bimodal when *L* = 1 with two nonzero modes. This usually occurs when *ρ*_0_ and *ρ*_1_ are both nonzero, which means that mRNA can be produced in all gene states. Clearly, increasing *L* results in a distribution that is more concentrated and has smaller noise, just as expected. Interestingly, the ways that the distribution is concentrated are significantly different for the two models. For the multi-OFF model, the left mode moves to the right and becomes higher as *L* increases, while the right mode remains almost unchanged and becomes lower. For the multi-ON model, however, the right mode moves to the left and becomes higher as *L* increases, while the left mode remains almost unchanged and becomes lower.

**Figure 5:**
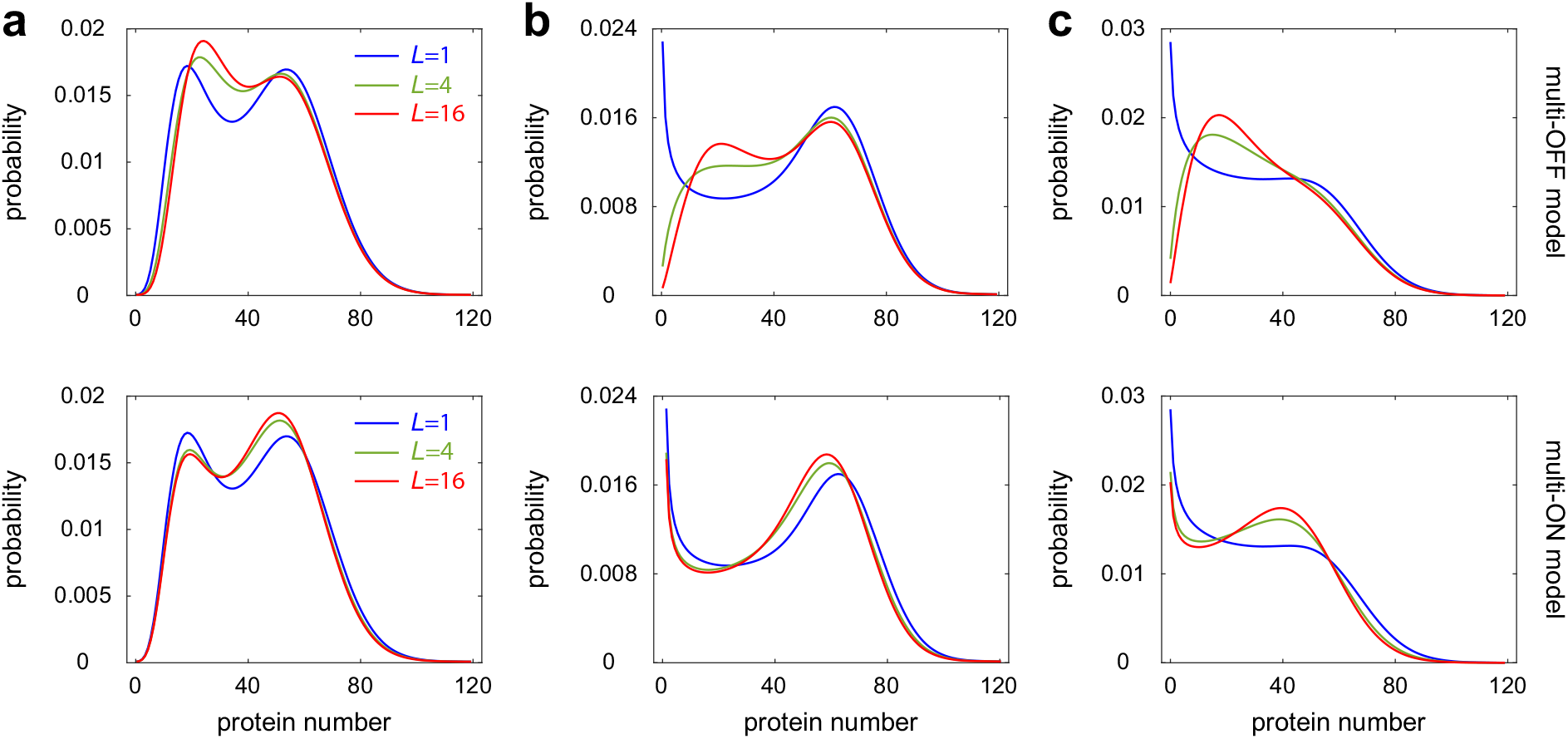
Influence of molecular memory on the steady-state distributions. (a) The case where the steady-state distribution is bimodal when *L* = 1 with two non-zero modes. The three curves show the distributions under different choices of *L*. The mean time staying at active states is 1/*u* while the mean time staying at inactive states is 1/*υ*. Other parameters are chosen as *u* = 0.35, *υ* = 0.46, *ρ*_0_ = 42, *ρ*_1_ = 10, *B* = 1.5 for the multi-OFF model and *u* = 0.35, *υ* = 0.46, *ρ*_1_ = 42, *ρ*_0_ = 10, *B* = 1.5 for the multi-ON model. (b) The case where the steady-state distribution is bimodal when *L* = 1 with a zero mode and a non-zero mode. The parameters are chosen as *u* = 0.4, *υ* = 0.7, *ρ*_0_ = 70, *ρ*_1_ = 0, *B* = 1 for the multi-OFF model and *u* = 0.4, *υ* = 0.7, *ρ*_1_ = 70, *ρ*_0_ = 0, *B* = 1 for the multi-ON model. (c) The case where the steady-state distribution is bimodal when *L* = 1 with an apparent zero mode and an inapparent non-zero mode. The parameters are chosen as *u* = 0.8, *v* = 0.8, *ρ*_0_ = 40, *ρ*_1_ = 0, *B* = 1.5 for the multi-OFF model and *u* = 0.8, *υ* = 0.8, *ρ*_1_ = 40, *ρ*_0_ = 0, *B* = 1.5 for the multi-ON model.

We emphasize that similar distribution properties also hold when the left mode is located at zero. In Fig. 5(b), the steady-state distribution is bimodal when *L* = 1 with a zero mode and a non-zero mode. This usually takes place when *ρ*_0_ = 0 or *ρ*_1_ = 0, which means that mRNA can be produced only in the active gene state. For the multi-OFF model, the zero mode moves rightwards as *L* increases and thus the distribution has two non-zero modes when *L* is large. Note that the fact that the multi-OFF model may produce two non-zero modes even when *ρ*_1_ = 0 has already been reported in [42]. For the multi-ON model, however, the zero mode still exists and the non-zero mode moves leftwards.

We next use these distribution properties to explain the counterintuitive enlargement for the TB and SB regions in the phase diagram for the multi-ON model. In Fig. 5(c), the steady-state distribution is bimodal when *L* = 1 with the left mode being dominant and the right mode being inapparent. This usually occurs around the cusp of the SB region in Fig. 4(b). For the multi-OFF model, the left mode moves to the right as *L* increases and thus U becomes more apparent when *L* is large. This is consistent with the observed shrinkage of the TB and SB regions in the phase diagram for the multi-OFF model. In contrast, for the multi-ON model, the right mode moves to the left and becomes higher with the increase of *L*. Since the left model is dominant, when *L* is large, the heights of the two modes become closer and bimodality becomes more apparent. This explains why the TB and SB regions enlarge in the phase diagram for the multi-ON model.

To summarize, we find that molecular memory has a marked impact on the dynamical behavior of the system when promoter switching is neither too fast nor too slow. Enhancing molecular memory in the inactive period reduces the TB and SB regions, while enhancing molecular memory in the active period enlarges them. Recent experiments have shown that in mammalian cells, the inactive periods for many genes has a non-exponential distribution [23, 24]. Our results suggest that this non-exponential behavior can significantly reduce transient and stationary bimodality, and thus stabilize both time-dependent and steady-state gene expression levels. This may explain why non-exponential inactive duration distributions are widely observed in nature.

## 6 Conclusions and discussion

In this paper, we derived the time-dependent distributions of mRNA and protein copy numbers for a multi-state gene expression model with complex promoter switching. Our model allows two different transcriptional rates, one corresponding to the transcriptionally active periods of the promoter and the other corresponding to the transcriptionally inactive periods. Moreover, our model takes into account two possibilities: (i) there are multiple inactive states of the promoter and only one active state; (ii) there are multiple active states and only one inactive state. The former corresponds to the case where the inactive period is non-exponentially distributed and thus has molecular memory, while the latter corresponds to the case where the active period has molecular memory. In each gene state, our model assumes that the gene product is produced in a non-bursty or bursty manner. The non-bursty case usually occurs for the mRNA dynamics while the bursty case usually occurs for the protein dynamics.

Note that our model assumes that all promoter states except state *G*_0_ have the same transcription rate. This is essential for a special function representation of the copy number distribution. Under this assumption, our model is equivalent to a non-Markovian two-state model with exponential switching one way and non-exponential switching the other way. While the non-Markovian formulation may be more natural, adding intermediate states allows us to describe the details of the underlying multi-step biochemical processes behind promoter switching. Recently, some papers [30, 31] have considered a general non-Markovian model with arbitrary switching time distributions between the active and inactive states. In such models, even in steady-state conditions, the special function representation of the copy number distribution can only be found for some very special cases.

Following previous work, we transformed the CMEs satisfied by the probability distribution into the PDEs satisfied by the generating function. The key to our analytical approach is to make nontrivial temporal, spatial, and functional transformations to obtain a generalized hypergeometric differential equation satisfied by the (transformed) generating function. Once this is done, the generating function can be represented as the linear combinations of fundamental solutions of the generalized hypergeometric differential equation. Moreover, we focused on the special cases where the promoter switching dynamics is very fast or very slow. By using multiscale simplification techniques of Markov chains, we obtained the simplified analytical expressions of the time-dependent distributions in the fast and slow switching limiting cases when the gene product number is initially zero and the promoter initially starts from the steady-state distribution of all gene states.

We then investigated the dynamical behavior of the multi-state model. We identified three dynamical phases of the multi-state model that are associated with three distinct types of time-evolution: unimodality, transient bimodality, and stationary bimodality. To understand the influence of molecular memory on the gene product number distribution, we determined the dynamical phase diagram of the multi-state model as the strength *L* of molecular memory varies. Based on the analytical solution and the phase diagram, we showed that (i) molecular memory has no influence on the time-dependent distribution when the promoter switches rapidly or slowly between all gene states; (ii) molecular memory significantly affects the time-dependent distribution when the promoter switching dynamics is neither too fast nor too slow. In the latter case, we found that molecular memory remarkably enhances the regions of transient bimodality and stationary bimodality if the active period has a non-exponential distribution, while the opposite is true if the inactive period is non-exponentially distributed.

For simplicity and analytical tractability, here we have not included any feedback regulation and cell cycle events. While we have considered an implicit description of gene product dilution due to cell division, via the effective gene product decay rate, it has recently been shown that in some parameter regimes, this type of model cannot capture the stochastic dynamics predicted by models with an explicit description of the cell cycle [62]. We hence anticipate that a more detailed gene expression model with the inclusion of feedback loops [17–21] and cell cycle events, such as cell growth, cell division, and gene replication [63–68] may display more complex behavior of time-evolution and may even introduce novel dynamical phases hitherto undescribed. These effects are currently under investigation.

## A Derivation of Eqs. (8) and (9) in the main text

Recall that in the main text, we have proved that

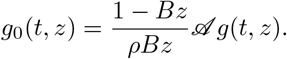

To proceed, we rewrite Eq. (6) in the main text in matrix form as

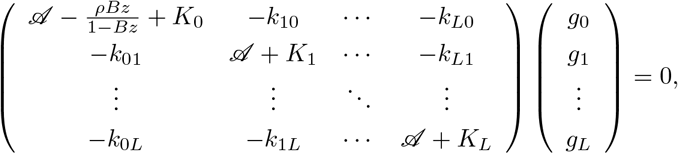

where 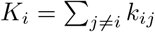. This equation, together with Eq. (7), shows that

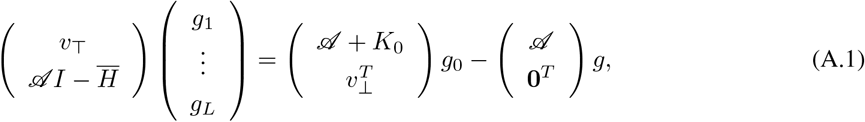

where *υ*_⊤_ = (*k*_10_, …, *k*_*L*0_), *υ*_⊥_ = (*k*_01_, …, *k*_0*L*_), *I* is the identity matrix, 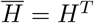 is the transpose of *H*, and 0 denotes the row vector whose components are all 0. Let *S* be an (*L* – 1) × *L* operator-valued matrix defined by

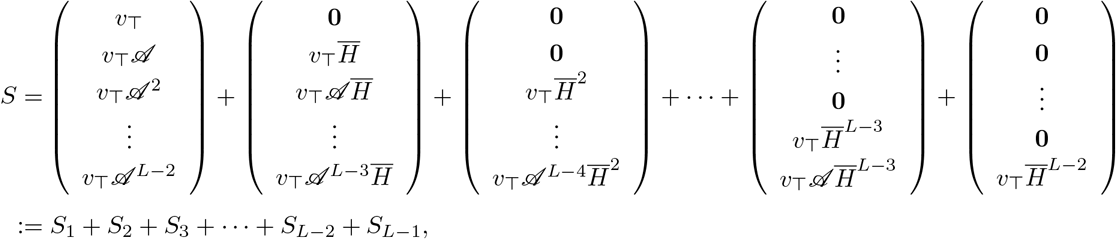

where the vectors 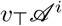 are defined by 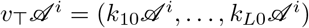. Then we have

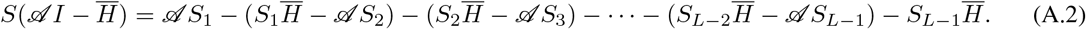

Note that

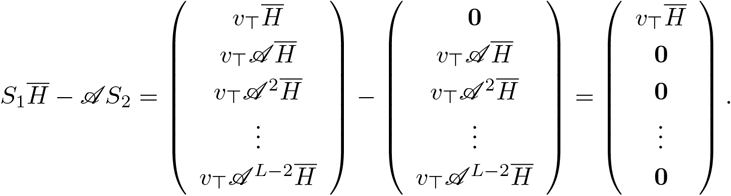

Similarly, we have

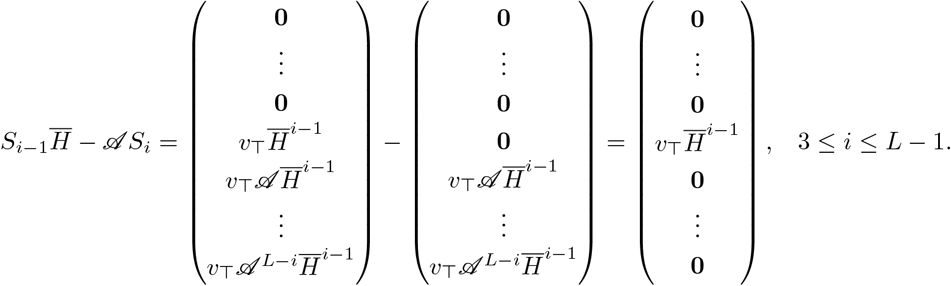

Inserting the above equations into Eq. (A.2), we obtain

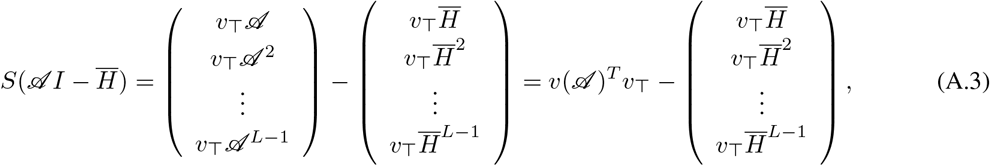

where 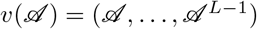. Next we define an *L* × (*L* + 1) operator-valued matrix *T* as

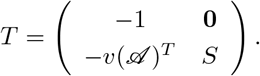

Then we obtain

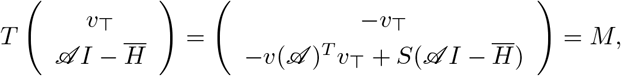

where

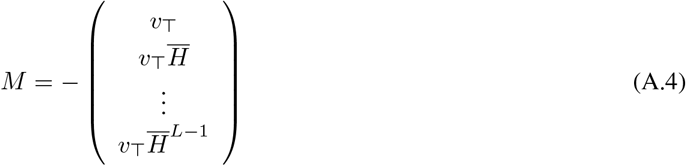

is a constant matrix which is called the Krylov matrix [69]. Multiplying *T* on both sides of Eq. (A.1) then yields

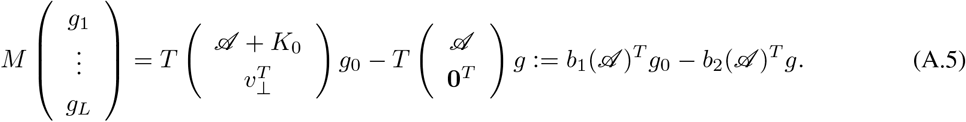

Since *g*_0_ can be expressed by *g* via Eq. (7), the above equation indicates that all *g*_1_, …, *g_L_* can also be expressed by *g*. It is clear that the matrix *T* only involves terms 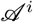 with *i* ≤ *L* – 1. Hence both 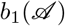 and 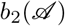 involve terms 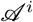 with *i* ≤ *L*. It then follows Eq. (7) that 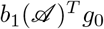 involves terms 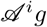 with *i* ≤ *L* + 1. This shows that 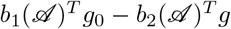 only involves terms 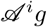 with *i ≤ L* + 1. To summarize, we obtain from Eq. (A.5) that

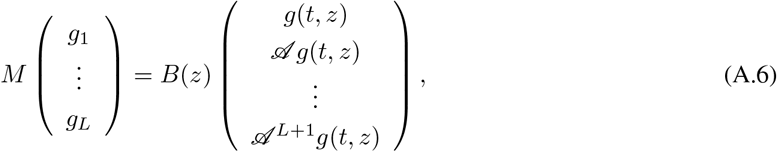

where *B*(*z*) is an *L* × (*L* + 2) matrix with its entries being functions of *z*.

In general, the matrix *M* defined in Eq. (A.4) may not be invertible. However, it is easy to check that *M* is always invertible when *L* = 1 and is always invertible when *L* = 2 for the multi-OFF and multi-ON models under all model parameters (see Appendix B for the proof). Even if *M* is not invertible, it can always be approximated by an invertible matrix with an arbitrary degree of accuracy. Thus, without loss of generality, we assume *M* to be invertible. By using the Cramer’s rule, we obtain

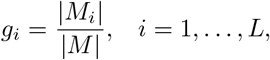

where |*M*| represents the determinant of *M*, and *M_i_* is the matrix formed by replacing the *i*th column of *M* by the vector 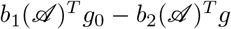. Since *g* = *g*_0_ + *g*_1_ + ⋯ + *g_L_*, we obtain

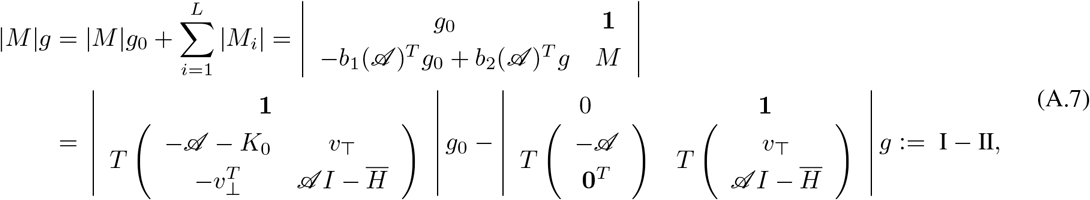

where 1 denotes the row vector whose components are all 1. We first analyze the first part I. Note that

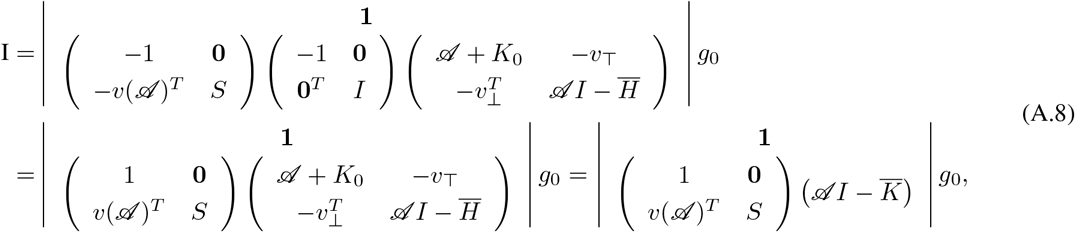

where 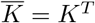. Since 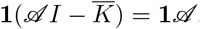, we have

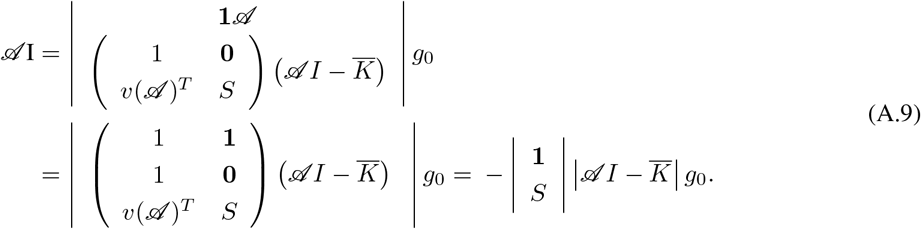

To proceed, note that

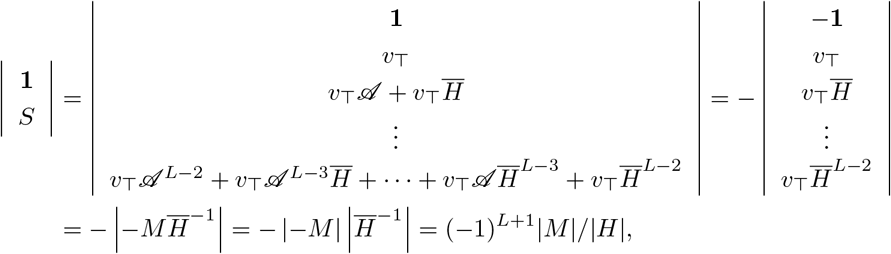

where we have used the fact that 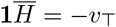. Since we have assumed that *M* is invertible, i.e. |*M*| ≠ 0, it follows from the above equation that |1^*T*^ *S^T^*| ≠ 0. Note that 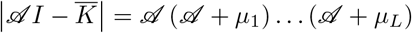. Combining this equation with Eqs. (A.8) and (A.9), and noting that the operator 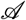 appears on both sides of the equation, we obtain

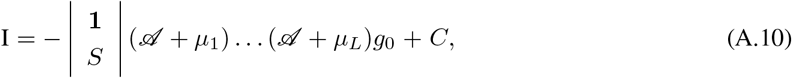

for some constant *C*. Finally, from the definition of I in Eq. (A.8), it is easy to see that *C* = 0.

We then analyze the second part II. Since 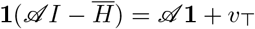, we have

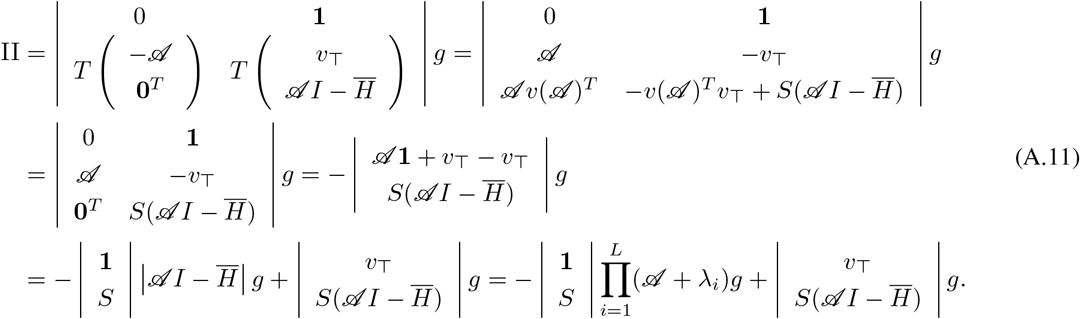

It is easy to see from Eq. (A.3) that

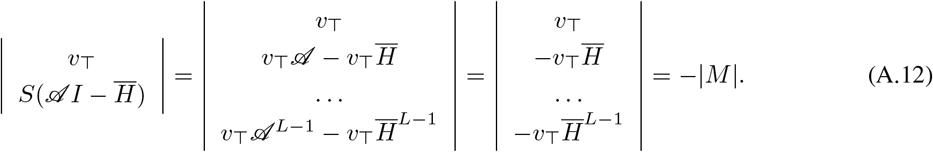

Inserting Eqs. (A.10), (A.11), (A.12) into Eq. (A.7) and using the fact that |1^*T*^ *S^T^*| ≠ 0, we obtain

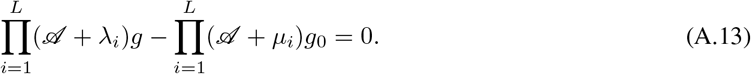

By induction and using Eq. (7), it is not difficult to prove that

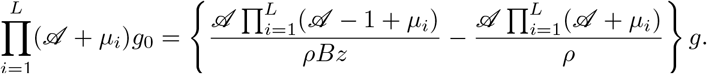

Inserting the above equation into Eq. (A.13) yields

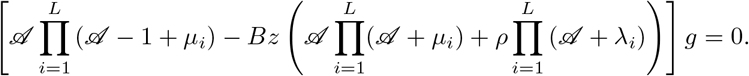

Note that

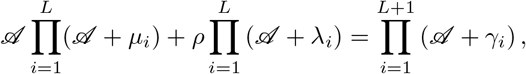

where *γ_i_* are determined by Eq. (10) in the main text. We then obtain Eq. (9) in the main text, from which it is clear that we can express 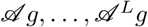 by 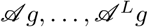. Combining Eqs. (7) and (A.6), we finally obtain Eq. (8) in the main text.

## B Time-dependent solution for the three-state model

For the three-state model with *L* = 2, we have

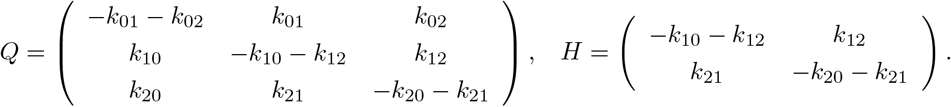

It follows from Eqs. (13) and (16) in the main text that

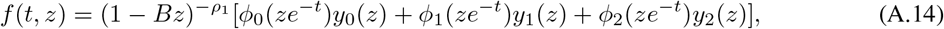

where

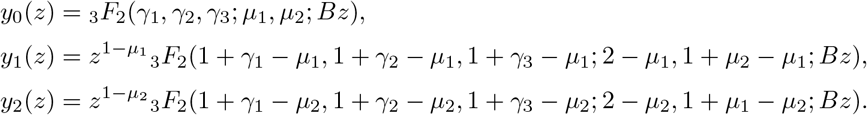

We next compute the explicit expressions of *ϕ_i_*(*z*). It is easy to check that *μ_i_*, **λ*_i_*, and *γ_i_* are determined by the following equations:

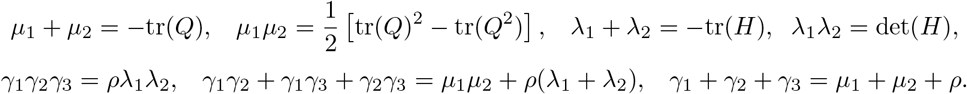

Straight forward computations show that Eq. (A.5) has the form of

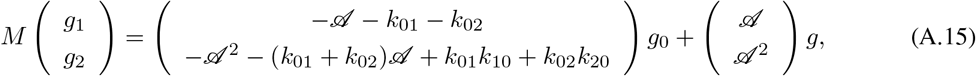

where

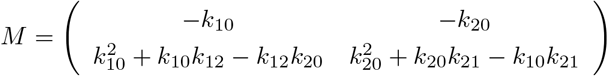

is a constant matrix. It is easy to check that

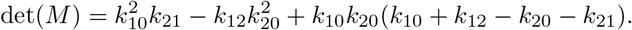

For the multi-ON and multi-OFF models, we have *k*_02_ = *k*_21_ = *k*_10_ = 0 and thus 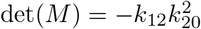, which is always non-zero. Since the analytical solution in the general case is too complicated, we next only focus on the multi-ON and multi-OFF models. From Eq. (7), it is not difficult to prove that

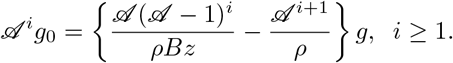

Applying this equation, Eq. (A.15) becomes

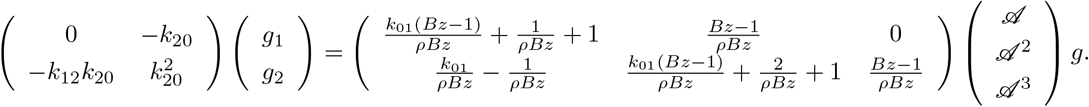

This shows that

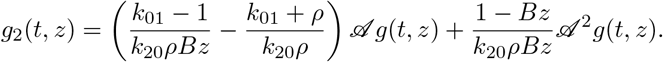

This equation, together with Eq. (7), yields

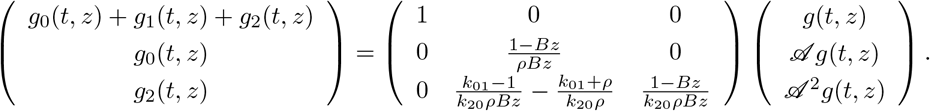

Applying Eq. (14) in the main text to the above equation and then setting *t* = 0, we obtain

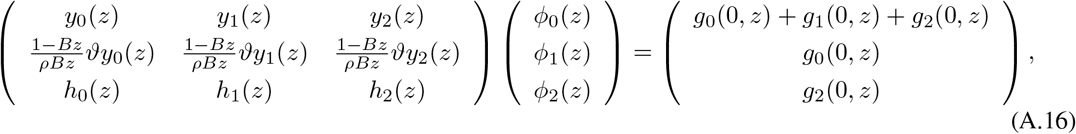

where

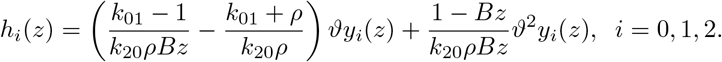

We then use Cramer’s rule to solve the above system of linear equations. Let *A*(*z*) denote the coefficient matrix of the above system of linear equations. It is easy to see that

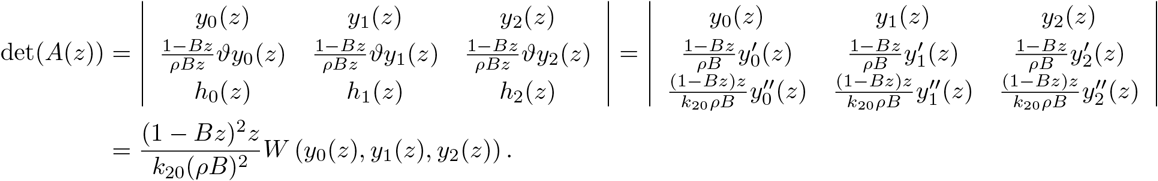

where *W* (*y*_0_(*z*), *y*_1_(*z*), *y*_2_(*z*)) is the Wronskian of the functions *y*_0_(*z*), *y*_1_(*z*), and *y*_2_(*z*). We emphasize here that the Wronskian is defined with respect to the original derivative, instead of *ϑ*. The following lemma gives the analytical expression for the Wronskian.

### Lemma 1.

Let *y*_0_(*z*), *y*_1_(*z*), …, *y_L_*(*z*) be the functions given in Eq. (12) in the main text. Then for each *L* ≥ 1, the Wronskian of these functions is given by

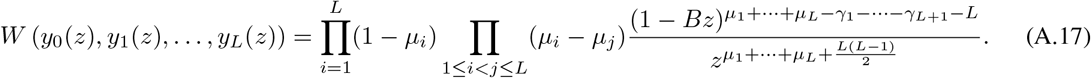

*Proof*. It has been shown in [70] that

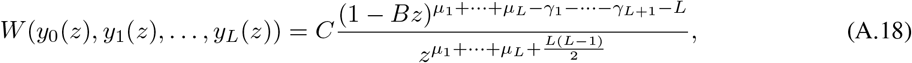

where *C* is an undetermined constant. Here we give the explicit expression of *C*. From Eq. (A.18), we have

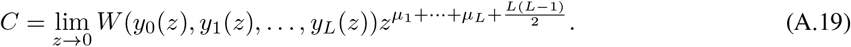

It is clear that *y*_0_(*z*) → 1 as *z* → 0. For any *n* ≥ 1, it is easy to check that 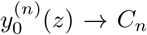 as *z* → 0 for some constant *C_n_*. For any *i* ≥ 1 we write 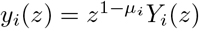, where *Y_i_*(*z*) is a generalized hypergeometric function which satisfies *Y_i_*(*z*) → 1 as *z* → 0. Note that

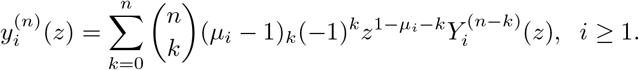

Thus for each *n* ≥ 0, we have

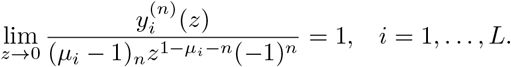

Using the above limit, we obtain

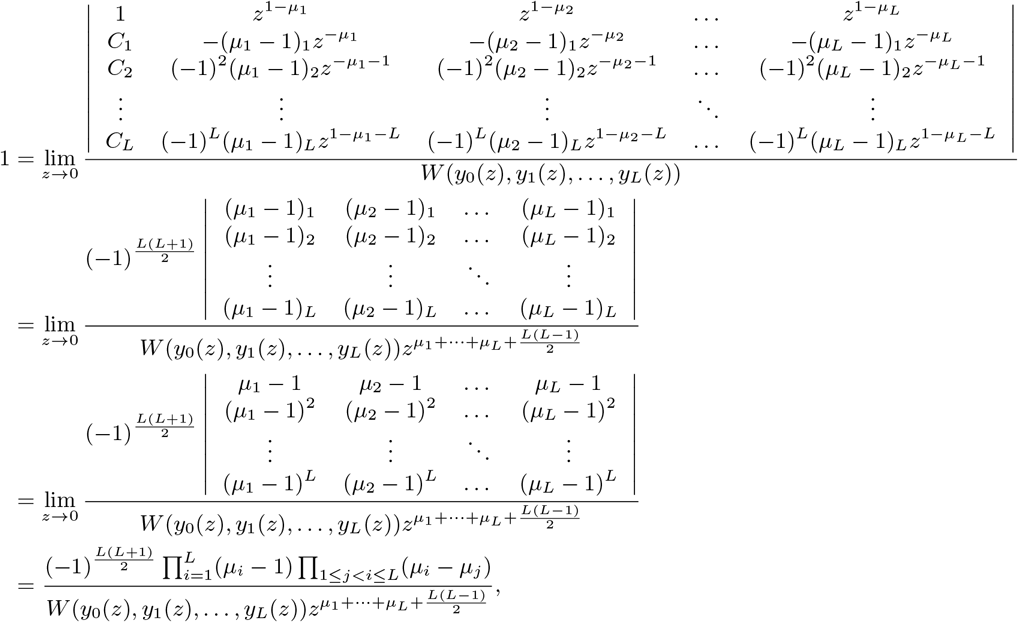

where the second equality follows from the expansion of the determinant with respect to the first column, the third equality follows from the fact that 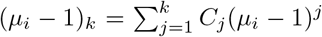 for some constants *C_j_* that are independent of *i*, and the fourth equality follows from the expression of the Vandermonde determinant. Combining the above equation with Eq. (A.19) yields

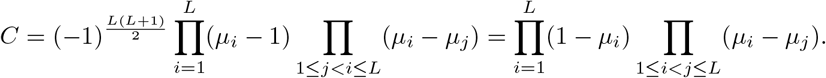

Inserting the above equation into Eq. (A.18), we finally obtain Eq. (A.17).

By using Lemma 1, we immediately see that for *L* = 2,

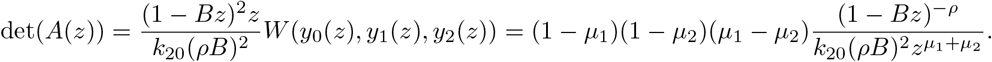

If the initial protein number is zero and the promoter is in state *G*_0_, then the initial conditions are given by

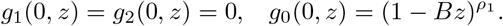

It then follows from Eq. (A.16) and Cramer’s rule that

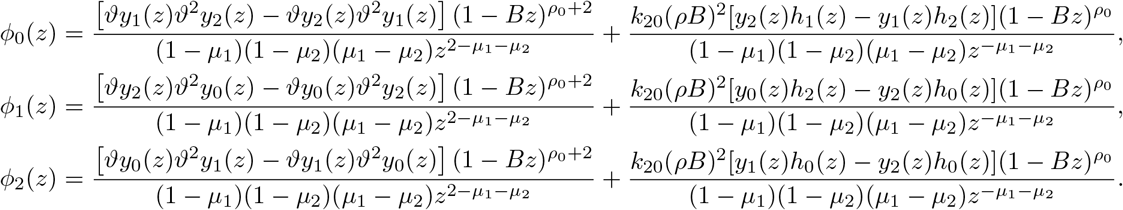

Inserting these equations into Eq. (A.14), we obtain the explicit expression of the generating function *f*(*t*, *z*).

## Acknowledgements

We are also grateful to Prof. Ramon Grima for stimulating discussions and Prof. Jiajun Zhang for valuable suggestions. We also thank Prof. Feng Jiao for sharing us with some important references. C. J. acknowledges support from the NSAF grant in National Natural Science Foundation of China with grant No. U1930402.

