## Supplementary Material 1 for "Analytical time-dependent distributions for gene expression models with complex promoter switching mechanisms"

### Contents

**1 Derivation of Eq. (19) in the main text** **2**

### 1 Derivation of Eq. (19) in the main text

By the definition of elementary symmetric polynomials and Eq. (10) in the main text, we have

$$\begin{aligned}\prod_{k=1}^{L+1}(x-\gamma_k) &= \sum_{k=0}^{L+1}\sigma_k(\gamma_1,\ldots,\gamma_{L+1})(-1)^k x^{L+1-k} = x^{L+1}+\sum_{k=1}^{L+1}\sigma_k(\gamma_1,\ldots,\gamma_{L+1})(-1)^k x^{L+1-k}\\ &= x^{L+1}+\sum_{k=1}^{L+1}\left[\sigma_k(\mu_1,\ldots,\mu_L)+\rho\sigma_{k-1}(\lambda_1,\ldots,\lambda_L)\right](-1)^k x^{L+1-k}\\ &= x^{L+1}+\sum_{k=1}^{L}\sigma_k(\mu_1,\ldots,\mu_L)(-1)^k x^{L+1-k}-\rho\sum_{k=0}^{L}\sigma_k(\lambda_1,\ldots,\lambda_L)(-1)^k x^{L-k},\end{aligned}$$

where we have used $\sigma_{L+1}(\mu_1,\ldots,\mu_L)=0$. Let $f$ be a function defined by

$$f(x)=\frac{x^{L+1}+\sum_{k=1}^{L}\sigma_k(\mu_1,\ldots,\mu_L)(-1)^k x^{L+1-k}}{x^L+\sum_{k=1}^{L}\sigma_k(\lambda_1,\ldots,\lambda_L)(-1)^k x^{L-k}}-\rho.$$

Clearly, the function $f$ has exactly $L+1$ zeros $\gamma_1,\ldots,\gamma_{L+1}$ in the complex plane. Moreover, it is easy to see that

$$\begin{aligned}f(x) &= x-\rho+\frac{\sum_{k=1}^{L}\left[\sigma_k(\mu_1,\ldots,\mu_L)-\sigma_k(\lambda_1,\ldots,\lambda_L)\right](-1)^k x^{L+1-k}}{x^L+\sum_{k=1}^{L}\sigma_k(\lambda_1,\ldots,\lambda_L)(-1)^k x^{L-k}}\\ &= x-\rho+\frac{\sum_{k=0}^{L-1}\left[\sigma_{k+1}(\mu_1,\ldots,\mu_L)-\sigma_{k+1}(\lambda_1,\ldots,\lambda_L)\right](-1)^{k+1} x^{L-k}}{\sum_{k=0}^{L}\sigma_k(\lambda_1,\ldots,\lambda_L)(-1)^k x^{L-k}} := x-\rho+h(x).\end{aligned}$$

Since the numerator and denominator of $h(x)$ are polynomials of the same degree, we have

$$\lim_{|x|\to\infty}h(x)<\infty.$$

Hence there exists constants $x_0>0$ and $C>0$ such that $|h(x)|\le C$ whenever $|x|\ge x_0$. Thus for any $\epsilon>0$, whenever $\rho\ge\max\{C/\epsilon, x_0/(1-\epsilon)\}$, we have

$$\begin{aligned}f\left((1+\epsilon)\rho)\right) &= \epsilon\rho+h\left((1+\epsilon)\rho\right)\ge\epsilon\rho-C\ge 0,\\ f\left((1-\epsilon)\rho)\right) &= -\epsilon\rho+h\left((1-\epsilon)\rho\right)\le-\epsilon\rho+C\le 0.\end{aligned}$$

By the mean value theorem, there exists $x_1\in[(1-\epsilon)\rho,(1+\epsilon)\rho]$ such that $f(x_1)=0$. Since $x_1$ is a zero of $f$, for convenience, we denote it by $\gamma_{L+1}$. By the arbitrariness of $\epsilon$, we have

$$\lim_{\rho\to\infty}\frac{\gamma_{L+1}}{\rho}=1.$$

Along the same line, we can also prove that the above limit holds when $\rho\to-\infty$. Finally we obtain

$$\lim_{|\rho|\to\infty}\frac{\gamma_{L+1}}{\rho}=1. \tag{1}$$

We then prove by induction on $k$ that

$$\lim_{|\rho|\to\infty}\sigma_k(\gamma_1,\ldots,\gamma_L)=\sigma_k(\lambda_1,\ldots,\lambda_L), \tag{2}$$

for any $k=1,\ldots,L$. To this end, we first prove that Eq. (2) holds for $k=L$. Recall that Eq. 10 in the main text is given by

$$\sigma_k(\gamma_1,\ldots,\gamma_{L+1})=\sigma_k(\mu_1,\ldots,\mu_L)+\rho\sigma_{k-1}(\lambda_1,\ldots,\lambda_L),\quad k=1,\ldots,L+1. \tag{3}$$

Setting $k=L+1$ in Eq. (3), we obtain

$$\prod_{i=1}^{L+1}\gamma_i=\rho\prod_{i=1}^{L}\lambda_i.$$

This equation, together with Eq. (1), shows that Eq. (2) holds for $k = L$. Then we prove the desired result by induction. Suppose that Eq. (2) holds for some $k = 2, \ldots, L$. By the definition of elementary symmetric polynomials, it is easy to see that

$$\sigma_k(\gamma_1, \ldots, \gamma_L, \gamma_{L+1}) = \sigma_k(\gamma_1, \ldots, \gamma_L) + \gamma_{L+1}\sigma_{k-1}(\gamma_1, \ldots, \gamma_L).$$

Then it follows from Eq. (3) that

$$\sigma_k(\gamma_1, \ldots, \gamma_L) + \gamma_{L+1}\sigma_{k-1}(\gamma_1, \ldots, \gamma_L) = \sigma_k(\mu_1, \ldots, \mu_L) + \rho\sigma_{k-1}(\lambda_1, \ldots, \lambda_L).$$

Since Eq. (2) holds for $k$, it follows from Eq. (1) that

$$\lim_{|\rho|\to\infty} \sigma_{k-1}(\gamma_1, \ldots, \gamma_L) = \sigma_{k-1}(\lambda_1, \ldots, \lambda_L),$$

which shows that Eq. (2) holds for $k-1$. Therefore, Eq. (2) holds for any $k = 1, \ldots, L$. Finally, by the fundamental theory of algebra, we obtain from Eq. (2) that

$$\lim_{|\rho|\to\infty} \gamma_i = \lambda_i, \quad i = 1, \ldots, L.$$
